# Cell-type-specific parallel pathways in the canonical cortical microcircuit

**DOI:** 10.64898/2026.04.23.720412

**Authors:** Chi Zhang, Casey M Schneider-Mizell, Bethanny P Danskin, Rachael Swanstrom, Erika Neace, Emily Joyce, Benjamin D Pedigo, Forrest C Collman, Nuno Maçarico da Costa

## Abstract

Information processing in the cortex depends on the integration of bottom-up and top-down signals through recurrent microcircuits spanning layers. Although the canonical microcircuit provides a framework for this integration, how these interactions are implemented at synapse resolution remains unclear. Here, we use large-volume electron microscopy reconstructions of mouse primary visual cortex to map the intralaminar and interlaminar connectivity of intratelencephalic (IT) neurons in layers 2/3 and 5. We find that layer 2/3 IT neurons formed a depth-dependent gradient of recurrent connectivity, with superficial (L2) and deeper (L3) neurons potentially forming two channels associated with top-down and bottom-up processing, respectively. These channels are preserved across layers via cell-type-specific pathways involving distinct L5 IT types, rather than collapsing into a single integrative pool. Moreover, each channel is regulated by a largely separate cohort of inhibitory interneurons, stabilizing recurrent excitation while limiting crosstalk. Together, these results reveal parallel, cell-type-specific processing streams embedded within the canonical circuit.

## Introduction

Cortical computation relies on the integration of bottom-up input with top-down contextual information originating from higher-order brain regions. Classical models of cortical organization attribute this integration to recurrent interactions among excitatory and inhibitory neurons distributed across cortical layers. A central framework for this view was formalized by Douglas and Martin, who proposed a canonical cortical microcircuit based on intracellular recordings^1^. In this model, recurrently connected excitatory neurons in supragranular (layers 2/3) and infragranular (layers 5/6) layers interact via interlaminar loops and shared inhibitory circuitry, with layer 4 and layer 2/3 serving as the principal recipient of feedforward thalamic input. Although this schematic necessarily simplifies the dense and heterogeneous cortical “thicket” first described by Ramón y Cajal, it established the view that cortical processing is recurrent and structured across layers.

Subsequent work not only explored the detailed anatomy of this microcircuit^2–5^, has shown that cortical microcircuits are shaped not only by feedforward thalamic signals but also by extensive feedback from higher-order cortical and subcortical areas. In the mouse primary visual cortex (VISp), neurons in layer 2/3 collectively act as a major integration hub, receiving bottom-up input from layer 4 and thalamus alongside top-down input from higher visual and associative areas^6^. Increasing evidence suggests that this supragranular population is itself heterogeneous: depth within layer 2/3 correlates with molecular identity, morphology, electrophysiological properties, and functional responses^7–11^. Upper layer 2/3 intratelencephalic (L2/3 IT) neurons receive stronger input from higher-order cortical areas, whereas deeper L2/3 IT neurons are more strongly driven by dorsal lateral geniculate nucleus (dLGN) and layer 4^7,12,13^. These depth-specific IT populations can be molecularly targeted ^10^ and exhibit distinct prediction-error responses^14,15^, consistent with biased integration of contextual and sensory information.

Comparable organizational principles extend beyond the visual system. In the primary somatosensory cortex, higher-order thalamocortical inputs preferentially target broad-tufted layer 2 IT neurons, whereas slender-tufted layer 3 IT neurons receive stronger first-order thalamic and layer 4 input^16,17^. Together, these findings suggest that parallel processing channels may coexist within supragranular layers, each biased toward integrating information from different hierarchical sources. However, how such channels are implemented within the recurrent cortical microcircuit—and whether they are preserved or collapsed as signals propagate across layers—remains unclear.

Layer 5 intratelencephalic (L5 IT) neurons provide a natural locus to address these questions, as they mediate interactions between supragranular processing and cortical output^18^. These neurons receive both bottom-up and top-down input ^12,19^ and form recurrent connections with L2/3 IT populations^20–23^. A recent functional study implicates L5 IT neurons in predictive processing in VISp^15^, yet how distinct L5 IT cell types interface with depth-specific supragranular circuits—and whether they act as generic or channel-specific integrators —remains largely unresolved. This question is particularly pressing given the growing recognition of L5 IT heterogeneity, with multiple transcriptomic, morphological, and functional subclasses identified in mouse cortex^8,24^.

In parallel, inhibitory circuitry adds an additional layer of complexity. Cortex contains a rich diversity of inhibitory neuron types distributed across layers^3,8,25–27^, and inhibitory neurons play a central role in stabilizing recurrent excitation, shaping temporal dynamics, and regulating information flow^28^. Although specific excitatory-inhibitory motifs have been described within VISp ^3,29–31^, it remains unclear whether inhibition acts as a global balancing force, mediates competition across processing streams, or is organized in a channel-specific manner that preserves parallel pathways.

Resolving these questions requires comprehensive, cell-type-resolved maps of synaptic connectivity across cortical layers. Large-volume serial-section transmission electron microscopy (ssTEM) now makes it possible to interrogate cortical microcircuits at synaptic resolution across many thousands of neurons. Here, we leverage the MICrONS connectomic dataset of mouse VISp^32^, which spans the cortex from pia to white matter with segmentation of neurons and synapses. To date, this dataset has been used primarily to dissect inhibitory circuitry^3,33^, whereas excitatory networks^31^ have remained comparatively less explored until now due to the limited availability of large-scale proofreading. Focusing on excitatory neurons in layers 2/3 and 5, we reconstruct their intralaminar and interlaminar recurrent connectivity, as well as associated disynaptic inhibitory pathways. By systematically mapping these excitatory–excitatory and excitatory–inhibitory networks, we aim to clarify how parallel processing channels are embedded within the cortical microcircuit and how high-resolution connectivity data refine and extend the classical concept of the canonical cortical microcircuit.

## Results

### VISp cell types in the MICrONS dataset

To analyze the circuitry, we build on the excitatory and inhibitory classification described in a previous study^3^. Transcriptomic and functional studies have shown that VISp L2/3 IT neurons can be classified into three molecularly defined types—*Cdh13/Trpc6/Chrm2*^9^ or *Adamts2/Agmat/Rrad*^8,10,34^—which are enriched in the upper, middle, and lower portions of L2/3, respectively. The expression of their marker genes follows continuous yet distinct spatial gradients across the depth of layer 2/3. Consistent with this molecular landscape, a connectomics classification^3^ based on morphological and synaptic features identified five excitatory IT types in layer 2/3, each occupying a relatively narrow depth range, but with partially overlapping domains. L2a is located at the superficial end, and L3b is positioned at the border of layer 2/3 and layer 4 (Fig. 1b). Furthermore, these types exhibit different ranges of total input synapses (Extended Data Fig. 1a). Layer 5 comprises IT, ET, and near-projecting (NP) neurons that have distinct projection patterns across brain areas^8,35^. Although the EM volume does not permit comprehensive reconstruction of long-range projections for definitive typing, local morphological and synaptic features provide sufficient criteria for classification. ET neurons are characterized by large somata, thick apical dendrites with prominent tufts, and numerous oblique branches, whereas L5 IT neurons display smaller somata, thinner apical dendrites, fewer oblique dendrites, and lower input synapse density^3,31,35,36^. Two types are identified in the L5 IT group: L5a IT neurons have more extensively branched tufts and higher input density compared to L5b IT neurons. L5a neurons are predominantly located in the upper portion of layer 5, whereas L5b neurons span the full depth of the layer. NP neurons are distinguished by their unique dendritic morphology and relatively small input synapse sizes. The inhibitory neurons across layers are categorized into 4 subclasses according to their output synaptic patterns as described previously^3^. Perisomatic targeting cells (PTC) are mostly basket cells that primarily target soma or proximal dendrites; distal dendrite targeting cells (DTC) include Martinotti and non-Martinotti cells that target distal basal or apical dendrites; sparsely targeting cells (STC) are neurogliaform cells and L1 interneurons that make sparse multisynaptic connections; and inhibitory targeting cells (ITC) are putative VIP+ neurons that mainly target other inhibitory neurons (Fig. 1b).

**Figure 1.**
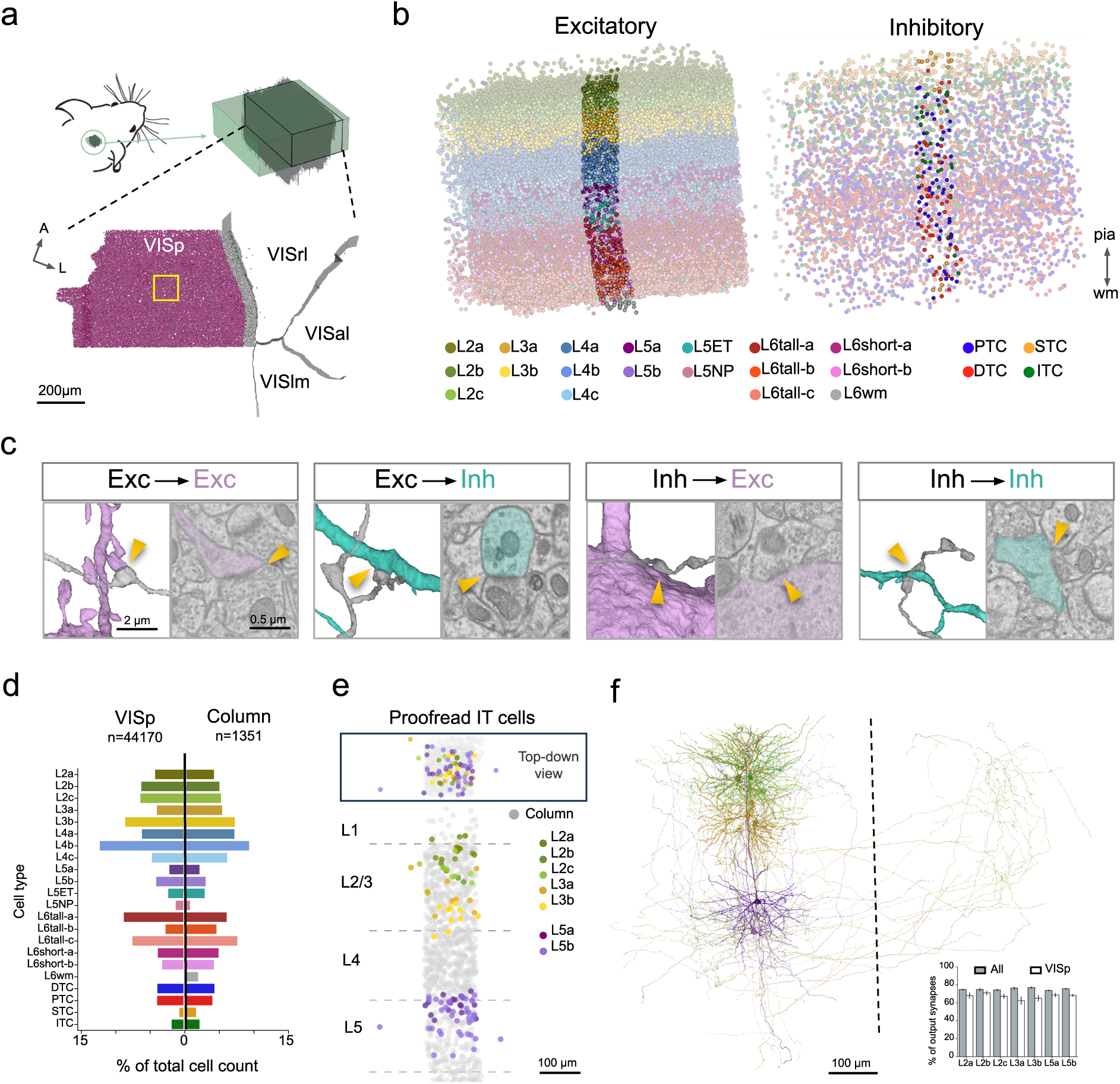
EM image volume and reconstruction of the mouse primary visual cortical circuits **a**. Illustration of the location and included cortical areas of the acquired EM volume. Solid circles indicate soma locations of primary visual cortical (VISp) neurons. Only neurons >50 µm from the areal border (magenta) were included for the analysis of connectivity in VISp, whereas those <50 µm (grey) were excluded. The yellow square indicates the column with extensive proofreading. **b**. Spatial distribution of excitatory (left) and inhibitory (right) VISp neurons across cortical layers (L), color-coded by predicted cell types. Neurons within the proofread column are highlighted. **c**. Representative meshes and raw micrographs of synapses (arrowheads) made among excitatory (Exc) and inhibitory (Inh) neurons, targeting different subcellular structures, including spines (1^st^ from left), shafts (2^nd^ and 4^th^), and soma (3^rd^). Postsynaptic structures are colored by cell types (Exc, magenta; Inh, teal). **d**. Percentages of cell types in VISp and proofread column. **e**. Top-town and side views of the distribution of exhaustively proofread intra-telencephalic (IT) cells. **f**. Representative meshes of layer 2/3 and layer 5 IT cells that demonstrate the dendritic and axonal morphology. The insert shows average percentages of output synapses with postsynaptic cell type identified for each IT type, across all the visual areas, or VISp area. VISal, anterolateral visual area; VISlm, lateralmedial visual area; VISrl, rostral lateral visual area; ET, extratelencephalic; NP, near-projecting; wm, white matter; PTC, perisomatic targeting cell; DTC, distal dendrite targeting cell; STC, sparse targeting cell; ITC, inhibitory targeting cell.

We focused our analysis on neurons whose cell bodies fell within a 100 × 100 µm wide column at the center of the VISp region (Fig. 1a,b). The column is representative of VISp in both cell type composition and their relative proportions (Fig. 1d) and contains the most thoroughly proofread neurons, with varied levels of axonal reconstruction (see Methods for proofreading details; Supplementary Information). Therefore, we selected L2/3 and L5 IT neurons located within and near the column (L2a, 8 cells; L2b, 9 cells; L2c, 9 cells; L3a, 9 cells; L3b, 10 cells; L5a, 17 cells; L5b, 38 cells) with comprehensive axonal reconstruction for connectivity mapping (Fig. 1e, f; see supplementary information for morphology). The cells were selected to cover the vertical and lateral distribution range of each cell type to avoid biased results. To ensure the area specificity, VISp neurons (n=44,170) at least 50µm away from the areal borders were included for VISp local connectivity analysis (Fig. 1a). The majority (75.5%) of the output synapses across cell types had postsynaptic partners with identified types with soma in the volume, and 67.7% on average were associated with VISp neurons (Fig. 1f). The analysis focused on microcircuits mediated by these fully proofread L2/3 and L5 IT neurons, involving either the column neurons or neurons across the entire VISp region.

### Depth-specific recurrent networks among L2 and L3 IT neurons

The canonical microcircuit emphasizes dense recurrent excitation within the supragranular layers. To examine this organization, we developed a cell-type-resolved view of recurrent excitation among layer 2/3 excitatory neurons, revealing how supragranular circuit architecture is further structured by neuronal depth and cell types. The excitatory cell types that occupied the superficial layer 2/3 (L2a and L2b IT neurons) both confined axonal arbors to layers 2/3 and 5 while largely avoiding layer 4. Compared to the two superficial types, the deeper L2/3 excitatory types (L2c, L3a, and L3b) extended their axons spanning layer 2 through layer 5, including layer 4 (Fig. 2a).

**Figure 2.**
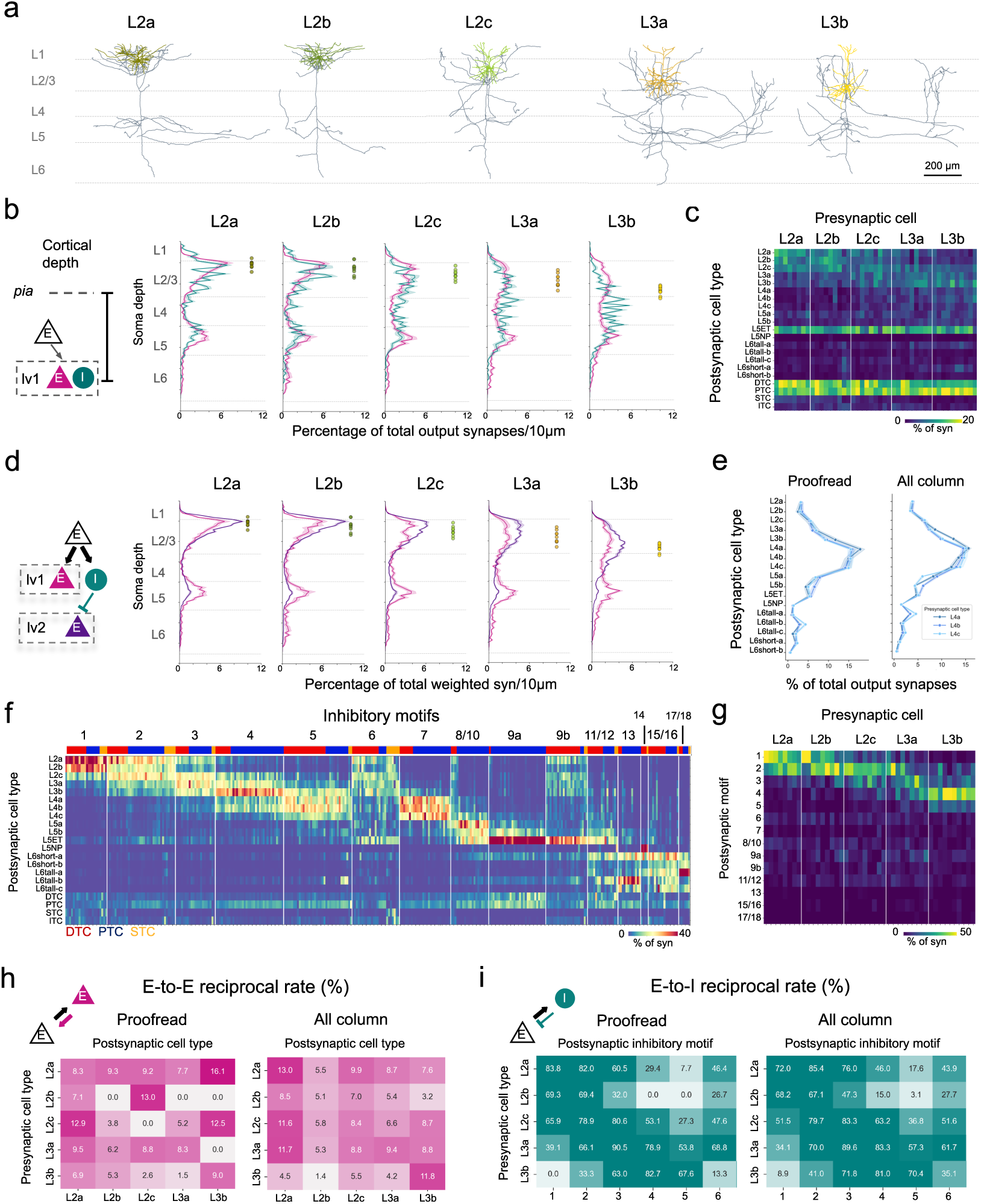
Depth-dependent recurrent connectivity of L2/3 IT neurons **a**. Representative skeletons of L2/3 IT neuron types. Dendrites are color-coded by neuron types. **b**. Cortical depth distributions of monosynaptic (lv1) excitatory (E, magenta) and inhibitory (I, teal) postsynaptic targets of L2/3 IT neurons. Each line indicates the mean ± SE of the percentage of total output synapses at 10µm intervals across cortical depth, averaged across starting L2/3 IT neurons. Soma locations of individual starting L2/3 IT neurons are indicated by color-coded filled circles on the right of each panel. **c**. Heatmap of the lv1 output connectivity of L2/3 IT neurons across different cell types. Cells are arranged by their types and soma depths. This arrangement applies to all the plots of individual cells. **d**. Cortical depth distributions of monosynaptic (lv1, magenta) and disynaptic (lv2, purple) excitatory postsynaptic targets of L2/3 IT neurons. **e**. Output connectivity of proofread (left) and all column (right) L4 IT neurons with VISp excitatory cell types. L4 IT types are color-coded. Each line indicates the mean ± SE of the percentage of total output synapses onto excitatory neurons. **f**. Heatmap of the output connectivity of each inhibitory motif. Top bar indicates the cell types of individual cells. **g**. Heatmap of the lv1 output connectivity of L2/3 IT neurons across different inhibitory motifs. **h**. Mean reciprocal rates between excitatory cell types for proofread (left) and all column (right) L2/3 IT neurons. **i**. Mean reciprocal rates between excitatory cell types and inhibitory motifs for proofread (left) and all column (right) L2/3 IT neurons.

We next examined whether the output connectivity also differed across L2/3 IT types by analyzing the laminar distribution and identities of their postsynaptic targets. In agreement with recurrent connectivity among L2/3 IT neurons detected by functional recordings^29,30,37–40^, all L2/3 IT types invested dense synaptic outputs within layers 2/3, with excitatory targets co-aligned to soma depth (Fig. 2b, Extended Data Fig. 1b). Given such alignment and depth-specific distribution of L2/3 IT neurons, each L2/3 IT type densely innervated its own and neighboring cell types in the vertical axis (Fig. 2c). In addition, L3 IT neurons on average made more synapses and higher percentage of their total output onto L4 IT neurons compared to L2 IT neurons (Extended Data Fig. 1c; L2 IT vs. L3 IT, n=26, 19 cells, 5.3±0.7 vs. 11.6±0.98 % of total output synapses, t-test, p<0.0001). Mutually, all three L4 IT types in the column preferentially targeted L3 over L2 IT neurons (Fig 2e; Extended Data. Fig. 1d,1e; 115 comprehensively proofread cells and 189 partially proofread cells). Furthermore, analyses of 30 comprehensively proofread putative dLGN axonal arbors in VISp also indicate that thalamocortical inputs preferentially target L3 IT neurons over L2 IT neurons (Extended Data Fig. 1f). These findings suggest that, consistently with established circuit architecture, L3 neurons were the predominant recipients of bottom-up feedforward inputs compared to L2 neurons.

Consistent with previous studies^41,42^, we found layer 2/3 excitatory neurons allocate a substantial fraction of their synapses to inhibitory neurons, which were typically located deeper than their excitatory targets (Fig. 2b, Extended Data Fig. 1b). The same correlation patterns were observed across all the column L2 and L3 IT neurons that had their axons reconstructed at least 100 μm from their soma (Extended Data Fig. 1b). Because of different depths between their excitatory and inhibitory targets, we wanted to know whether L2 and L3 IT neurons drive excitation and inhibition to influence separate downstream populations. To do this, we mapped the disynaptic pathways: first identifying inhibitory neurons directly contacted by L2 and L3 IT neurons (lv1, monosynaptic), and then identifying the excitatory neurons targeted by those inhibitory cells (lv2, disynaptic). In addition to the column inhibitory neurons, we proofread 214 additional inhibitory neurons whose soma were located within a 50µm halo surrounding the column (Extended Data Fig. 2a). These neurons span layer 1 through 5 and include all four inhibitory neuron subclasses. With a total of 413 proofread inhibitory neurons within and surrounding the column, we observed no differences in laminar distribution between the full population of inhibitory targets and the proofread subsets of inhibitory neurons postsynaptic to each IT type (Extended Data Fig. 2b). Therefore, the proofread inhibitory population provides a reliable and unbiased representation of the monosynaptic target population.

Monosynaptic and disynaptic connections onto excitatory neurons showed both convergence and laminar specialization. For each L2 and L3 IT type, the density peaks of monosynaptic and disynaptic excitatory targets in layer 2/3 were aligned (Fig. 2d, Extended Data Fig. 1b), indicating that excitation and inhibition converge onto the same depth-defined excitatory populations. However, although L2 and L3 IT neurons made similar amounts of monosynaptic connections onto excitatory and inhibitory neurons in layers 2/3 and 5, their disynaptic inhibitory effects were much more concentrated in layer 2/3 than the monosynaptic excitation (Fig. 2d, Extended Data Fig. 3). This pattern suggests that L2 and L3 IT neurons exert tight inhibitory control of supragranular networks, whereas their influence on layer 5 is conveyed primarily through excitation. We further found that both DTCs and PTCs contributed to L2 and L3 IT neuron-mediated inhibitory networks (Fig. 2c). Overall, we observed no differences between the depth distributions of the PTC and DTC targets across IT types in layer 2/3, or their disynaptic targets (Extended Data Fig. 3), although we did observe a systematic shift in how proximal L2 and L3 IT inputs were onto the soma of PTC versus DTC neurons (Extended Data Fig. 2c).

Overall, these results indicate that the layer 2/3 excitatory population is further organized into depth-specific networks. In this organization, L2 IT neurons form a recurrent circuit potentially biased toward top-down inputs, whereas L3 IT neurons participate in a circuit that preferentially integrates thalamic feedforward input^7,12,13^. This organization is consistent with previous functional studies in the mouse VISp^14,15^.

### L2 and L3 IT neurons receive prominent reciprocal feedback inhibition

Recurrent connectivity within layer 2/3 is a defining feature of the canonical cortical microcircuit and is often described at the level of cell types. To determine how prominently this connectivity pattern is expressed within layer 2/3 networks and to move beyond population-level descriptions, we next examined recurrent coupling at the level of individual neurons by systematically identifying reciprocal synaptic connections.

Across individual comprehensively proofread neurons, 8.1% of connected L2/3 IT pairs had reciprocal connectivity. Expanding this analysis to cover all column L2 and L3 IT neurons with at least partial proofreading yielded a nearly identical reciprocal rate of 8.3%. As is typical of excitatory-excitatory connections, >85% of these reciprocal connections contained only a single synapse (Extended Data Fig. 2e). Although some pairs of L2/3 IT types had higher rates than others, they were all below 15% (Fig. 2h). In contrast, L2/3 IT neurons organized markedly stronger reciprocal connections with inhibitory networks, with an average reciprocal rate of 42.6% between proofread L2/3 IT and proofread inhibitory neurons.

The high degree of inhibitory reciprocal connectivity supports prior functional findings, which indicate that interneurons highly interconnect with layer 2/3 excitatory neurons that share similar feature selectivity^43,44^. Importantly, this also suggests that inhibition within the layer 2/3 networks is largely organized within excitatory cell types, rather than across types, as it has been proposed^45^. To further test this idea, we leveraged our previous work identifying inhibitory motifs with strong cell-type specificity^3^ and asked whether excitatory neuron types preferentially target the same inhibitory motifs that, in turn, project back onto them^31^. Specifically, we categorized all the proofread inhibitory neurons, except ITC neurons, into motifs based on their output connectivity in VISp, enriching for neurons that preferentially target specific excitatory neuronal populations (Fig. 2f, Extended Data Fig. 2d; see supplementary materials for the morphology of all motifs). The clustering approach was similar to that previously applied to column inhibitory neurons^3^, and we reproduced comparable motifs with minor modifications (see Methods). Some motifs (e.g., 1-5 and 7) were layer-specific, whereas others (e.g., 6, 8/10, and 9) targeted distinct cell types with sparser input to other layers. As expected, L2 IT types preferentially recruited motifs 1 and 2, whereas L3 IT types engaged more motifs 3 and 4 that strongly target their own and neighboring types, often with multiple synapses in each connection (Fig. 2g, i; Extended Data Fig. 2f). L3 IT neurons, especially L3b, targeted motif 5 that inhibited L3b and L4 IT neurons, which is consistent with their disynaptic excitatory targets distribution (Fig. 2d). Together, these results indicate that the excitation among individual L2/3 IT neurons is predominantly unidirectional, whereas their interactions with inhibitory targets are more reciprocal. Therefore, the recurrent excitation within layer 2/3 operates mainly at the population level, and recurrent inhibition occurs at both population and single-cell levels.

### L2 and L3 IT neurons selectively target layer 5 extratelencephalic (ET) neurons

We next examined excitatory projections from L2/3 IT neurons to layer 5. These projections were initially described as constituting a predominantly feedforward pathway^46^. However, within the canonical microcircuit framework^1^, they have also been proposed to participate in recurrent interlaminar interactions that are central to several computational models^45,47^. Given the cellular diversity of layer 5, we sought to determine how inputs from supragranular IT neurons are distributed across distinct layer 5 cell types. In particular, layer 5 ET and IT neurons serve distinct functional roles, with ET neurons being the principal cortical output to subcortical targets, whereas IT neurons primarily mediate cortico-cortical communication^48^. All L2 and L3 IT types made significantly more synapses onto L5 ET than L5 IT neurons (Figs. 2c, 3a, Extended Data Fig. 1c; t-test, p<0.0001 for both L2 and L3 IT types). In addition, L2 and L3 IT populations showed subcellular compartment-specific targeting patterns onto L5 ET apical dendrites. L2 IT neurons strongly targeted the tufted arbors in layers 1 and 2, whereas L3 IT neurons allocated their synapses onto the apical dendrite stalk at the border of layers 3 and 4 (Fig. 3a). Although layer 5 contains more IT than ET neurons (70 IT vs. 39 ET in the column), individual ET neurons have more total input synapses than IT neurons in total (8,200 vs. 2,986 synapses/cell on average), as well as in each layer (Fig. 3b). Consequently, the chance to connect to ET neurons is about 1.5 times that of IT neurons, assuming all the synaptic sites are evenly distributed in space. To test whether this ET-biased targeting pattern was solely opportunity-based (Peters’ rule) or involved cell-type selectivity, we shuffled L2 and L3 IT output connections by randomly reassigning each L2 and L3 IT output synapse to one of nearby synapses that matched for postsynaptic E/I identity (see Methods). The observed true connectivity to L5 ET was higher than shuffled connectivity across layer 2/3 IT types, with true vs. shuffled ratios greater than 1 (Fig. 3c). In contrast, L2 IT neurons exhibited ratios below 1 when targeting L5b IT neurons, indicating selective avoidance of L5b type, a pattern not observed for L3 IT neurons. Furthermore, there were no differences between observed and shuffled connectivity onto L2 and L3 IT neurons (Fig. 3c), consistent with a space-dependent wiring mode within L2/3. Therefore, L2 and L3 IT neurons employ two local wiring strategies: proximity-governed connectivity with other L2 and L3 IT neurons and cell-type-selective targeting of L5 ET and IT neurons (Fig. 6a).

**Figure 3.**
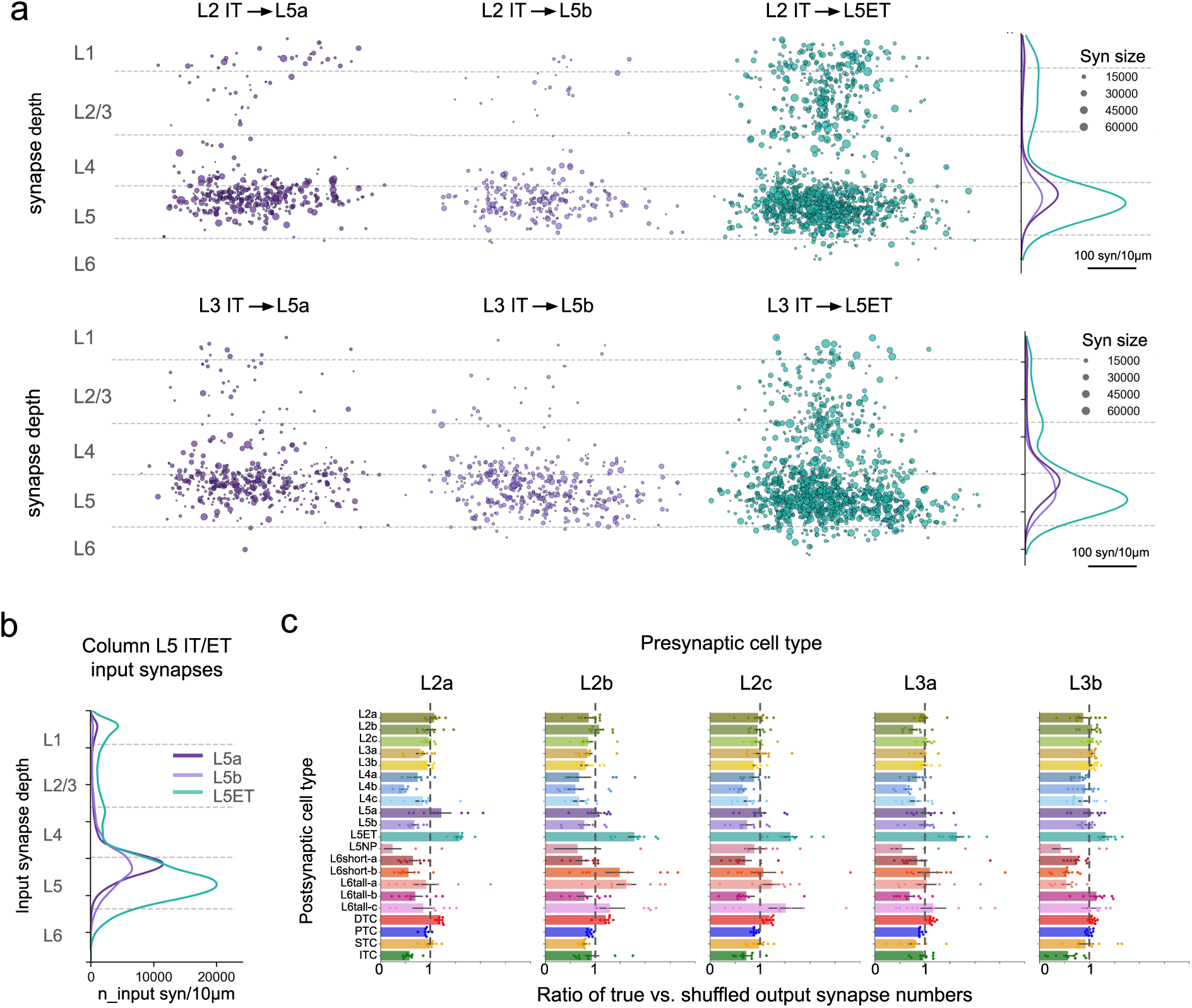
Cell type-selective connectivity of L2/3 IT neurons in layer 5 **a.** Scatter plot illustrating spatial locations and sizes of synapses across layers made by the proofread L2 (top) and L3 (bottom) IT neurons onto L5a IT, L5b IT, and ET neurons. KDE estimated depth distributions of synapses are shown on the right. **b.** KDE estimated depth distributions of all input synapses of L5 IT and ET neurons in the proofread column. **c.** Ratios of the true and shuffled output connectivity of each L2/3 IT cell type. Each bar graph shows mean ± SE ratios across cell types, with overlaid points representing ratios of individual cells. A dashed line indicates a ratio of 1 in each graph.

### Diversity of morphology and connectivity within the VISp L5 IT population

Although L5 IT neurons are secondary recipients of layer 2/3 IT output within layer 5, they nonetheless receive a substantial fraction of their total input from L2 and L3 IT neurons, and in turn serve as a prominent recurrent connection back to layer 2/3^22,23^.

Given the molecular and anatomical diversity of L5 IT neurons ^8,24^, we first sought to resolve distinct L5 IT cell types within the EM volume. Previous work on this dataset has already shown that L5 IT population can be separated into L5a and L5b IT neurons based on their dendritic features and synapse properties^3^. In this work, we added axonal reconstructions that elucidated this distinction further (Fig. 4a): L5a neurons typically extended ascending axons that sparsely branched in layer 1 and dense horizontal axons within layer 5, whereas L5b IT neurons were manually segregated into three groups based on their axonal projections to layer 2/3. The first two groups both formed dense axonal arbors in layer 2/3 rather than in layer 5, but with different ramification patterns (Fig. 4a). Axons of L5b-1 (n=13) predominantly branched in upper layer 2/3, whereas L5b-2 (n=16) axons spanned the full depth of layer 2/3. The third group (L5b-s, n =9) extended only sparse axon collaterals into layer 2/3 and was highly variable (Supplementary Materials). Consistent with their distinct axonal innervation in layer 2/3, L5a and L5b groups differed in the proportion of their synaptic outputs targeting layer 2/3 IT neurons (Extended Data Fig. 4a). All four L5 IT groups sent side branches into HVAs without a clear area-specific preference (Supplementary Information).

**Figure 4.**
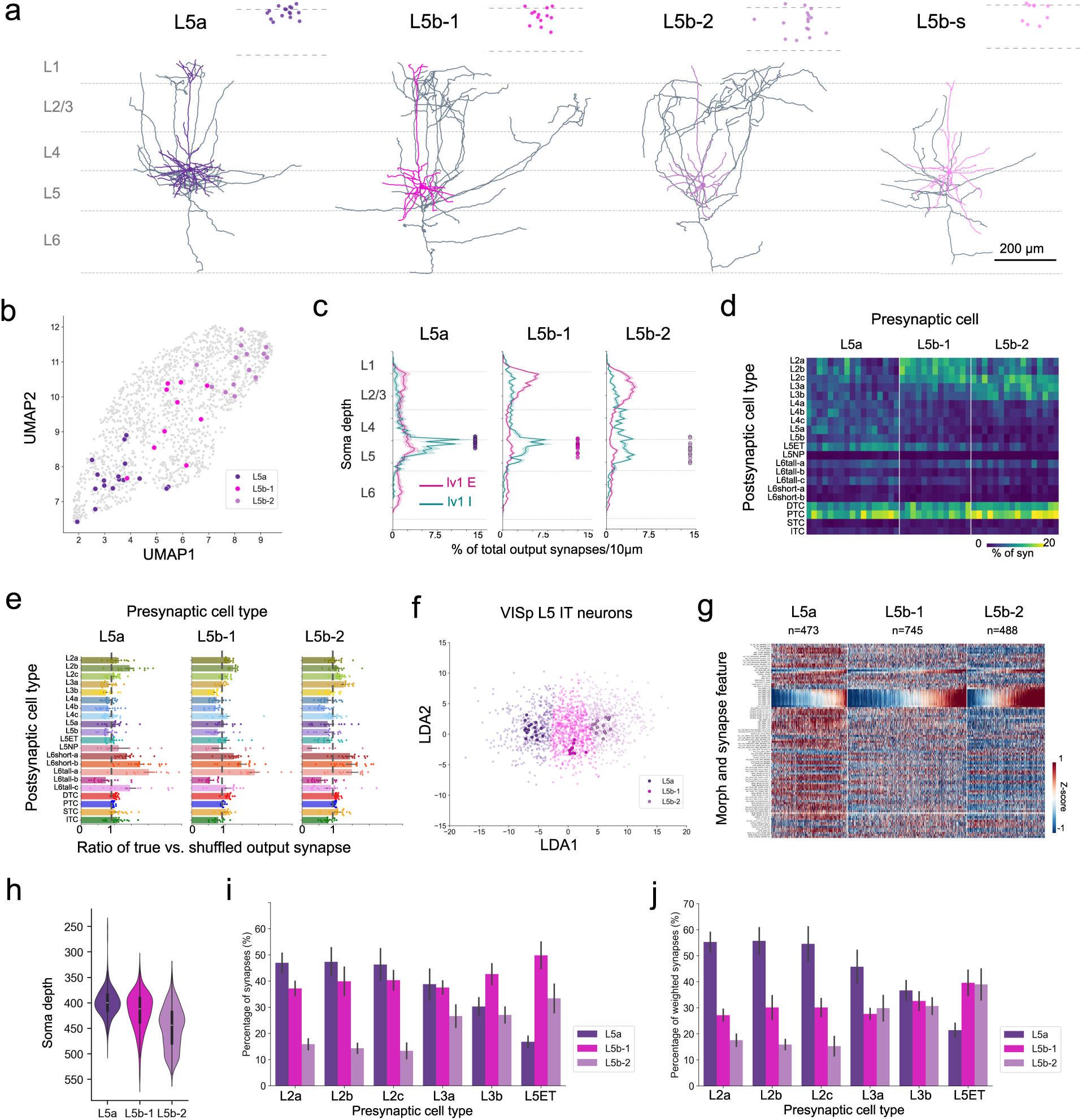
Cell type-specific connectivity between L5 and L2/3 IT neurons **a.** Representative skeletons of L5 IT neuron types. Soma locations of individual neurons in layer 5 are indicated for each cell type. Dendrites are color-coded by cell types. **b.** Uniform manifold approximation and projection (UMAP) embedding of synaptic and morphological features of VISp L5 IT neurons (grey) with proofread cells color-coded by cell types. **c**. Cortical depth distributions of monosynaptic (lv1) excitatory (E, magenta) and disynaptic (lv2) inhibitory (I, teal) postsynaptic targets of L5 IT neurons. Soma locations of individual starting L5 IT neurons are indicated by color-coded filled circles on the right of each panel. **d.** Heatmap of the monosynaptic output connectivity of L5 IT neurons across different cell types. **e**. Ratios of the true and shuffled output connectivity of each L5 IT neuron type. **f**. Linear discriminant analysis (LDA) clustering of VISp L5 IT neurons, trained using proofread L5 IT neurons as labels. Proofread cells are highlighted. **g**. Heatmap of z-scored features of individual VISp L5 IT neurons. **h**. Violin plot of L5 IT soma depth distribution. **i.** Output targeting patterns of L2/3 IT and L5 ET neurons onto L5 IT neuron types. Each set of three bars represents the distribution of postsynaptic L5 IT types for a given presynaptic type. **j.** Same as in (i), but with target proportions normalized by the abundance of each L5 IT type; for each presynaptic type, the proportions across the three L5 IT types sum to 1.

In addition to distinct axonal arborization, L5b groups also differ in other aspects. L5b-1 neurons exhibited sparsely branched tufted apical dendrites and horizontally extended basal dendrites, whereas most L5b-2 neurons possessed untufted apical dendrites and radially symmetric, star-shaped basal dendrites. The two groups also differed in laminar position: L5b-1 neurons occupied the upper half of layer 5, whereas L5b-2 neurons were distributed throughout the full thickness of layer 5 (Fig. 4a,c). Finally, L5b-1 neurons received substantially more total input synapses than L5b-2 neurons (Extended Data Fig. 4b; t-test, p<0.0001). To assess whether the observed differences reflect distinct cell types, we compiled anatomical and synaptic features (see Methods) of dendrites from 1706 L5 IT neurons in the dataset and performed an unsupervised embedding of these features (Fig. 4b). The resulting dendritic feature space appeared largely continuous, yet manually labeled L5b-1 and L5b-2 neurons occupied distinct regions (Fig. 4b), supporting their classification as different types despite graded feature variation. As expected, L5a neurons were also separated from both L5b populations in the feature space, consistent with their identity as a distinct L5 IT type. The L5b-s population, on the other hand, showed a non-specific distribution across the feature space occupied by L5b types. Possible explanations include that the sparse population includes extreme variants of L5b-1 and L5b-2 types, or insufficient features and limited sample size for reliable prediction, or that the group comprises mixed cell types, as suggested by the heterogeneous dendritic and axonal morphologies (Supplementary materials). With the sparsity of the L5b-s group and their low connectivity in layer 2/3, we focused on L5a, L5b-1, and L5b-2 neurons to compare their associated circuits within VISp to investigate their cell-type-specific roles in local activity modulation.

### L5 IT neurons form cell-type-specific channels with layer 2/3

After establishing the diversity of layer 5 IT cell types, we next examined their recurrent projections back to layer 2/3.

Unlike L5 ET neurons, whose local arbors innervated more inhibitory than excitatory neurons in VISp^31^, all three L5 IT types were biased towards excitatory neurons (Extended Data Fig. 4c). We first examined whether L5 IT types target excitatory and inhibitory cells in layer-specific patterns. Consistent with their distinct axonal morphology, L5a neurons had a larger fraction of layer 5 excitatory targets than L5b neurons; L5b neurons heavily innervated excitatory neurons at different depths of layer 2/3 (Fig. 4c), with L5b-1 targeting upper layer 2/3 and L5b-2 targeting middle and lower layer 2/3.

The output connectivity of each L5 IT type across excitatory cell types suggests specific circuits (Fig. 4d, Extended Data Fig. 4d). In layer 2/3, L5a neurons made sparse connections, with a few cells making more synapses onto L2 IT neurons, likely due to the branches around the border of layers 1 and 2. In comparison, L5b-1 and L5b-2 neurons substantially targeted different L2/3 IT populations, with L5b-1 biased toward L2 IT neurons and L5b-2 toward L3 IT neurons (Fig. 4d, Extended Data Fig. 4d; t-test, p = 0.002 for L2, p < 0.0001 for L3). In contrast to proximity-dependent targeting among L2/3 IT cells, a synapse shuffling test suggests that the sublayer-specific targeting by L5b-1 and L5b-2 is not random (Fig. 4e).

To investigate whether L2/3 IT neurons also have preferences for specific L5 IT types, we performed supervised dimensionality reduction for all the L5 IT neurons with their anatomical and synaptic features, using the proofread cells as training labels (Fig. 4f; see Methods). The analysis separated L5 IT neurons into three groups with distinct feature profiles and laminar distributions similar to the three labeled types (Fig. 4g,h). We then examined the connectivity of proofread L2/3 IT cells with these cell-type-identified L5 IT neurons. Notably, L2 and L3 IT neurons presented distinct preferences for L5a versus L5b-2, whereas no preference for L5b-1 (Fig. 4i). All three L2 IT types formed over three times more synapses onto L5a than onto L5b-2 neurons, consistent with the observed avoidance of L5b neurons by L2 IT neurons (Fig. 3c). In contrast, L3 IT neurons allocated a significantly greater proportion of their output to L5b-2 cells. This opposing preference became more pronounced after correcting for differences in the abundance of L5 IT types (Fig. 4j). This pattern is unlikely to reflect depth-dependent targeting, as L2 and L3 IT axons span the full depth of layer 5 (Fig. 2a).

Together, the excitatory connectivity between L2/3 and L5 IT neurons suggests the existence of two overlapping but distinct channels with similar architectural motifs. One channel is biased toward L2 IT neurons and is predicted to preferentially integrate top-down inputs, whereas the other is enriched for L3 IT neurons and is biased toward bottom-up sensory input. Each of these overlapping subcircuits preferentially routes excitation to distinct populations of L5 IT neurons (Fig. 6b).

### Layer 5 IT neurons couple excitation and inhibition across supragranular circuits

With the cell-type-specific excitatory channels between layers 2/3 and 5, we next examined how layer 5 neurons coordinate inhibitory interactions within and across these channels.

Although most inhibitory neurons targeted by L5 IT neurons had soma located in layer 5 (Fig. 4c), the monosynaptically and disynaptically innervated excitatory populations of each L5 IT type broadly followed similar distributions in layer 2/3 (Fig. 5a). For example, L5b-1 type preferentially drove inhibition onto L2 IT neurons, whereas L5b-2 type drove stronger inhibition onto L3 IT neurons (Extended Data Fig. 4f). Therefore, L5 IT neurons mediate both excitatory and inhibitory modulation onto L2/3 IT neurons in a cell-type-specific pattern (Fig. 6a).

**Figure 5.**
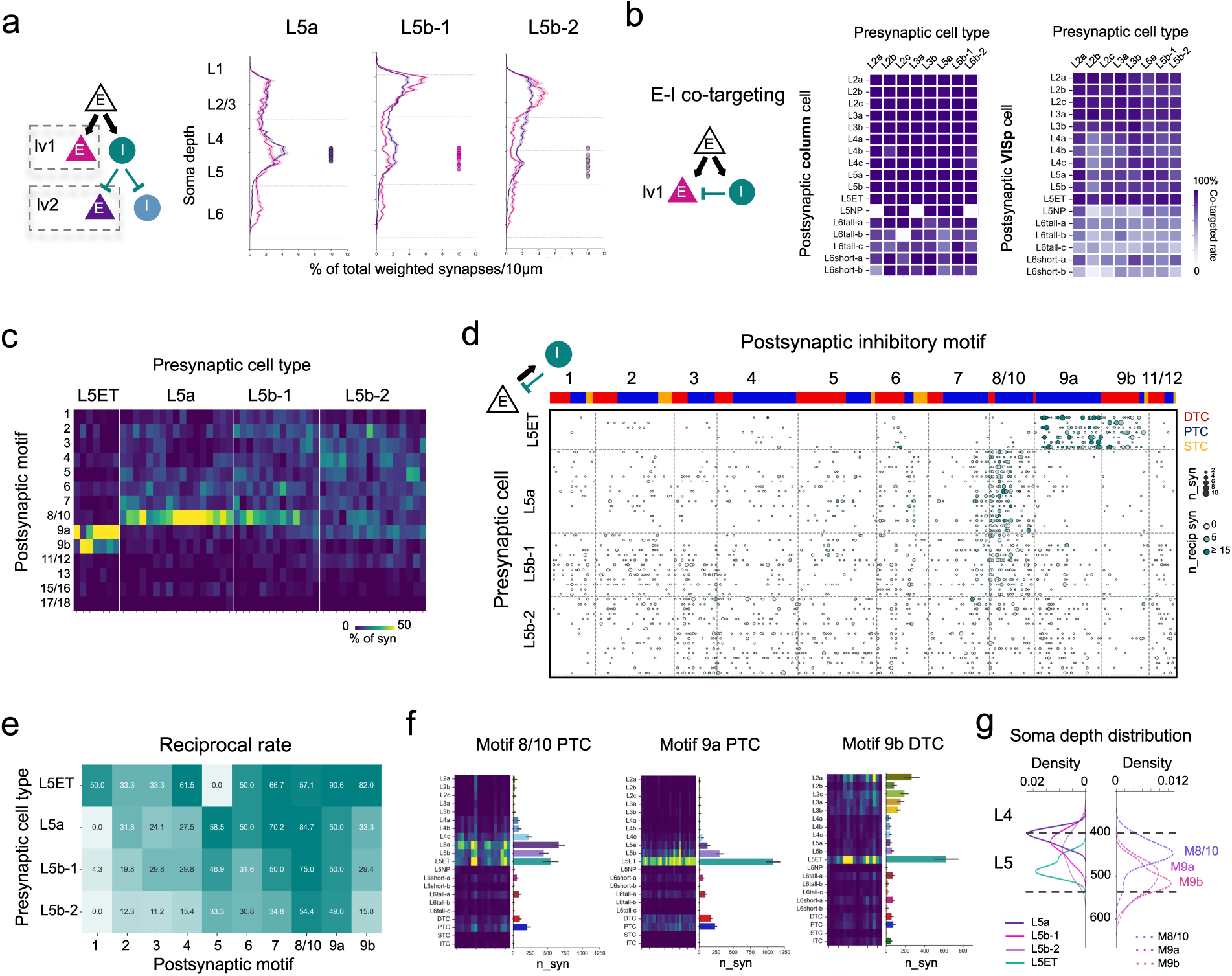
L5 IT and ET neurons drive distinct inhibitory circuits in layer 5 **a.** Cortical depth distributions of monosynaptic (lv1, magenta) and disynaptic (lv2, purple) excitatory postsynaptic targets of L5 IT neurons. **b**. Heatmap showing the average co-targeted rate of each excitatory cell type by monosynaptic excitation and disynaptic inhibition from L2/3 and L5 IT neurons. The left panel includes lv1 column excitatory targets, and the right panel includes all lv1 VISp excitatory targets. **c.** Heatmap of the lv1 output connectivity of L5 ET and IT neurons across different inhibitory motifs. **d.** Connectivity matrix of individual pairs of presynaptic proofread L5 ET/IT neurons and postsynaptic proofread inhibitory neurons arranged by motifs and cell types (top bar). Circle size indicates the synapse number of forward (pre-to-post) connections. Circle hue indicates the number of synapses of reciprocal connections. **e.** Reciprocal rates of L5 ET and IT neuron types with inhibitory motifs. **f.** Output connectivity of Motif 8/10 and 9a PTC, indicated by output synapse number of individual cells (left) and averaged output synapse number (right). **g.** Soma depth distribution of L5 ET and IT neurons (solid lines), and three inhibitory motifs (dashed lines), heavily innervated by L5 excitatory neurons.

**Figure 6.**
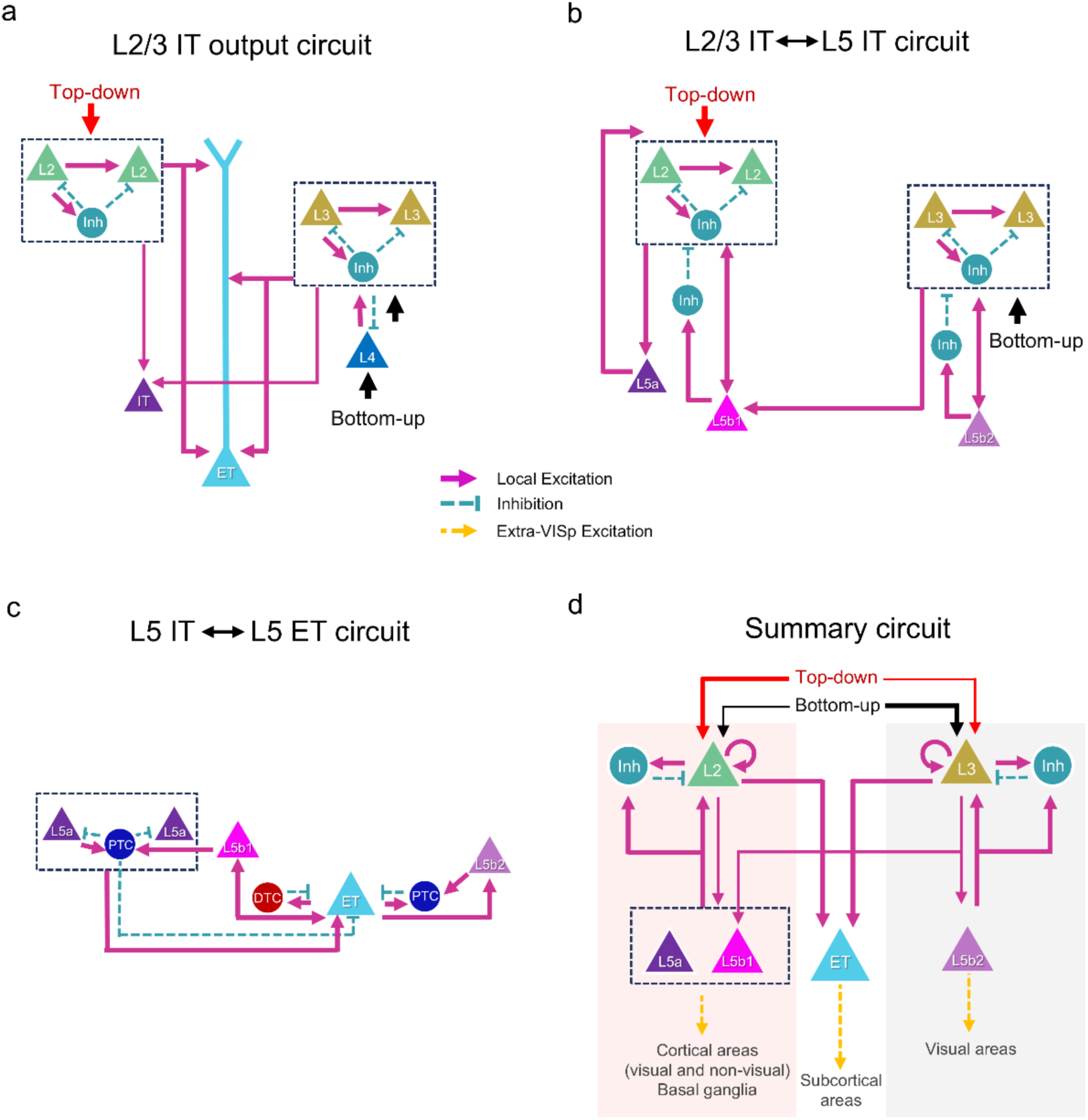
VISp L2/3 and L5 IT type-specific circuitry. **a.** Recurrent networks of L2 and L3 IT neurons in layers 2/3 and their output connectivity with L5 excitatory neurons. **b**. Cell-type-specific channels between L2/3 and L5 IT neurons. **c.** Cell-type-specific excitatory and inhibitory circuits mediated by L5 IT and ET neurons. **d.** Summary diagram of cell-type-specific local and long-range channels participating in top-down and bottom-up information processing. Only connections related to critical cell type specificity are indicated in this diagram. The extra-VISp projection patterns are estimated by comparing our IT neurons with L5 IT neurons reported in Sorensen et al. 2026 ^24^ and Gao et al. 2026 ^52.^

This organization raised the question of whether individual L2/3 IT neurons were co-targeted by excitation and inhibition originating from the same L5 IT neurons - a connectivity pattern we refer to as monosynaptic excitation and disynaptic inhibition (MEDI) and mimics the well-described feedforward inhibition^49^. Despite the distance between layers 2/3 and 5, 100% of the monosynaptic column L2/3 IT targets of each L5 IT type were co-targeted (Fig. 5b). This high co-targeted rate extended to most cell types except those in layer 6, where we observed direct excitation but not disynaptic inhibition from L5 IT neurons (Fig. 5a). The excitation-dominant innervation in layer 6 is unlikely to result from incomplete inclusion of inhibitory neurons, as the distributions of all versus proofread-only inhibitory targets of L5 IT neurons were indistinguishable (Extend Data Fig. 4e). When we expanded the analysis to all monosynaptic VISp targets, including those postsynaptic to unproofread monosynaptic inhibitory neurons not captured in the disynaptic group, co-targeted rates for L2/3 IT, L5 IT, and ET neurons remained above 75% (Fig. 5b). Similar patterns of MEDI were also observed for L2/3 IT driven circuits, suggesting that coordinated excitation and inhibition onto shared targets is a common motif in VISp local circuitry. Together, beyond their morphological distinctions, L5 IT types exhibited highly differentiated output connectivity and formed cell-type-specific pathways with L2/3 IT neurons through tightly coupled excitatory and inhibitory microcircuits.

### Cell-type-specific interaction between L5 IT and ET neurons

A final component of information flow within the excitatory network is recurrent excitation among layer 5 cell types.

As two major cell classes in layer 5, L5 IT and ET neurons are known for substantially interconnecting each other^31^. In our analysis, we observed distinct connectivity with ET neurons by different L5 IT types. L5a neurons had the strongest output onto L5 IT and ET cells compared to L5b neurons (Fig. 4d, Extended Data Fig. 4f), whereas L5b-2 neurons made minimal connections with layer 5 excitatory neurons. In contrast, analysis of comprehensively proofread column L5 ET neurons (n=7; Supplemental Information)^31^ revealed a preferential targeting of L5b over L5a neurons, without selectivity among L5b types (Fig. 4i,j).

### Divergent inhibitory motif recruitment by IT and ET neurons

We next assessed the inhibitory circuits driven by L5 IT and ET neurons. Here, we observed a striking segregation of inhibitory circuits driven by IT and ET cell subclasses. The three layer 5 inhibitory motif groups (8/10, 9a, 9b) were recruited in a cell-type-specific manner: L5 IT neurons preferentially engaged motif 8/10, which is enriched in PTCs, whereas L5 ET neurons predominantly recruited motifs 9a (PTC-enriched) and 9b (DTC-enriched) (Fig. 5c)^31^. Consistent with this segregation, ET and IT populations showed minimal co-targeting of inhibitory neurons across subclasses (Extended Data Fig. 4g), indicating largely parallel inhibitory architectures.

Different L5 IT types exhibited an additional layer of cell-type-specificity in their connectivity with these motifs. L5a neurons strongly targeted motif 8/10, which preferentially inhibited L5 IT and ET neurons and often formed reciprocal feedback onto L5a cells (Fig. 5d-f, Extended Data Fig. 4f). In contrast, L5b-1 showed weaker targeting of this motif, and L5b-2 only sparse, low-reciprocity connections. Importantly, this targeting gradient is more cell-type-selective than depth-dependent, as neurons of different L5 IT types located at similar depths still exhibited distinct connectivity profiles (Fig. 5d,g). ET neurons strongly targeted motifs 9a and 9b, both of which selectively innervated ET neurons with high reciprocity (Fig. 5c-f). In contrast, L5 IT neurons showed limited engagement: both L5b IT types sparsely but consistently targeted motif 9a, whereas L5a neurons did so only sporadically (Fig. 5d). Despite occupying similar spatial locations with motif 9a (Fig. 5g, Extended Data Fig. 2d), motif 9b neurons were rarely targeted by L5 IT neurons (Fig. 5d). Such avoidance of the DTC-enriched motif by L5 IT neurons is consistent with their weaker DTC-mediated disynaptic inhibition in layer 5 compared to PTC-mediated inhibition (Extended Data Fig. 4e). Similar to the motif 8/10-targeting pattern, these distinct connectivity with motifs 9a and 9b cannot be explained by depth-dependency, as no correlation was observed between soma position and motif targeting across individual L5 IT neurons.

Together, these findings demonstrate that inhibitory circuitry in layer 5 is organized into cell-type-specific, largely non-overlapping networks aligned with distinct excitatory output classes. Rather than participating in a shared inhibitory pool, IT and ET neurons engage dedicated inhibitory motifs that form reciprocal, closed-loop circuits (Fig. 6c). This organization suggests that inhibition in layer 5 is tailored to the functional roles of different excitatory cell types, supporting distinct modes of cortical computation and output.

### A wiring diagram of cortical excitation

The analyses above converge on an integrated excitatory wiring diagram for mouse primary visual cortex, summarized in Figure 6, which links depth-dependent supragranular circuits to cell-type-specific translaminar pathways. At the core of this organization is a graded but structured separation of excitatory processing streams within layer 2/3, coupled to distinct IT neuron populations in layer 5.

Within supragranular layers (Fig. 6a), excitatory connectivity is organized primarily by depth and spatial overlap. L2 and L3 IT neurons form dense recurrent networks with neighboring excitatory populations, and their mutual connectivity can largely be explained by the alignment of dendritic and axonal arbors. This depth-dependent organization gives rise to partially overlapping but biased supragranular pathways.

These supragranular pathways are not collapsed as signals propagate to deeper layers. Instead, they are selectively routed through layer 5 via cell-type-specific connectivity with distinct L5 IT types (Fig. 6b). L5a IT neurons preferentially receive inputs from L2 IT neurons, whereas L5b-2 IT neurons are more strongly targeted by L3 IT neurons. Recurrent projections from L5 IT neurons back to layer 2/3 further reinforce this organization, forming cell type-specific interlaminar loops. L5b-1 neurons occupy an intermediate position, receiving input from both supragranular populations and projecting broadly, consistent with a more integrative role. Downstream convergence occurs primarily at ET neurons, which receive input from multiple IT pathways (Fig. 6c).

Taken together, these findings reveal a wiring diagram in which excitatory cortical circuits are organized as overlapping, cell-type-specific pathways rather than as a single recurrent pool. Spatial proximity governs recurrent connectivity within layers, whereas cell-type-specific targeting governs interlaminar routing. This architecture provides a structural basis for parallel processing of bottom-up and top-down signals within the canonical cortical microcircuit (Fig. 6d).

## Discussion

### The diversity of cell types

Morphology and connectivity are classical and complementary criteria for defining neuronal cell types^50^. For IT cells in layer 2/3, we adopted the cell-type definitions established in the MICrONS dataset^3,32^. The depth-dependent connectivity patterns we observe are consistent with the known continuum of morphological and gene-expression variation across layer 2/3 excitatory types^7,8,11,24,51^. For layer 5 IT types, we extended the classification previously presented in the MICrONS dataset ^3^. We identified three types of L5 IT neurons in the mouse VISp EM volume, distinguished by their dendritic and axonal morphology, synaptic properties, and connectivity. Such diversity is consistent with findings from recent studies using other modalities. Transcriptomic analyses have classified VISp L5 IT neurons into 5 molecular types, with different depth distributions^8,34^. A multimodal study by Sorensen and colleagues identified four L5 IT types based on combined morphological, physiological, and molecular features^24^. Comparisons of soma depth and morphology suggest that our L5a, L5b-1, and L5b-2 types most closely correspond to the L4/5 IT, L5IT-1, and L5IT-2 groups reported by Sorensen et al., respectively^24^ (Extended Data Fig. 5). Many L4/5 IT neurons have denser axons in L1 than our L5a cells. This discrepancy may reflect incomplete reconstruction, as ascending axons from L5a neurons often extend laterally toward the edge of the image volume, and the MICrONs reconstructions lack the top 10-15 μm of L1. The correspondence between L5b-1 and L5IT-1 is less clear than L5b-2 and L5IT-2, due to the absence of axonal morphology of L5IT-1. It was also difficult to identify cells in our dataset corresponding to their fourth group, L5IT-3 Pld5, due to the small number of examples and the lack of axonal reconstructions. Nevertheless, from a large-scale anatomical study by Gao et al. that reconstructed >18,000 mouse cortical neurons^52^, we found VISp L5 IT neurons with dendritic and axonal morphology that match our three L5 IT types (Extended Data Fig. 5).

### Multiple channels across the cortical microcircuit

By leveraging cell-type classifications and comprehensive axonal proofreading, we were able to systematically chart the wiring diagram of IT neurons, providing a circuit-level framework for understanding how cortical networks integrate bottom-up sensory evidence with top-down contextual signals. Numerous theoretical frameworks, including predictive processing and hierarchical inference models^45,47,53,54^, posit the existence of parallel feedforward and feedback streams. Our findings provide synaptic-level evidence that such streams are implemented as overlapping but distinct microcircuits defined by neuronal depth in layer 2/3 and cell-type-specific connectivity with L5 IT neurons.

Consistent with known molecular, morphological, and electrophysiological gradients in L2/3 IT neurons^7–11^, we find that depth-dependent organization within supragranular layers gives rise to biased circuit architectures. L2 IT neurons are spatially positioned to preferentially integrate top-down inputs, whereas L3 IT neurons overlap with feedforward thalamic drive. Our study provides direct anatomical evidence that L2 and L3 IT neurons that occupy the upper and lower layer 2/3 differentially contribute to partially segregated microcircuits. L3 IT neurons make significantly more synaptic connections with L4 neurons, both pre- and postsynaptically, than L2 IT neurons. L2 IT neurons more densely innervate the tufted apical dendrites of L5 ET neurons, a critical subcellular compartment for feedback integration, whereas L3 IT neurons target the apical dendrite shaft, a hotspot for feedforward inputs^12,55^. This organization is consistent with known differences in the inhibitory populations targeting L 2 and L 3 IT neurons^3^. In particular, L3 and L4 IT neurons receive shared inhibition from specific inhibitory cell types, while L2 IT neurons receive inhibitory input from a different population of neurons. Such structured inhibitory targeting provides an additional mechanism by which L2- and L3-centered circuits can remain functionally differentiated despite their spatial overlap.

These biases correlate with the depth of dendritic and axonal arbors within layer 2/3: for example, L2c neurons located in mid-L2/3 form more synapses with layer 4 than L2a/b neurons, whereas L3b neurons at the bottom of L2/3 exhibit the strongest interaction with layer 4. Similarly, recurrent connectivity between L2 and L3 IT neurons can largely be predicted by the spatial overlap of their dendritic and axonal arbors.

In contrast, connectivity of L2/3 IT neurons with layer 5 neurons, as well as with some inhibitory cell types^3^, cannot be explained by spatial overlap alone. Instead, we find that these connections are formed with strong cell-type specificity. Together, these observations indicate that L2/3 IT neurons recruit both depth-dependent and depth-independent cell-type-specific wiring principles to assemble microcircuits within and across cortical layers. As a result, L2 and L3 IT neurons form distinct circuit motifs. These are not strictly segregated channels, as direct connections between L2 and L3 IT neurons are present; nevertheless, consistent biases in connectivity are evident.

### Layer 5 IT cell types as channel-specific rather than generic integrators

Bottom-up- and top-down-biased supragranular channels are not confined to layer 2/3 but remain coordinated through cell-type-specific recurrent loops involving layer 5 IT neurons. L5 IT neurons occupy a pivotal position in cortical hierarchies, receiving extensive input from superficial layers while projecting broadly to other cortical areas. Indeed, L5 IT activities are found to sharpen feature selectivity in L2/3 in the primary auditory cortex^56^. In VISp, L5 IT neurons provide ‘teaching signals’, that are different from bottom-up inputs relayed from L4, to L2/3 IT neurons to perform prediction^15^. Although they have often been conceptualized as generic integrative hubs that pool supragranular activity, accumulating functional evidence suggests a more selective role in shaping cortical representations. Our findings provide a circuit-level basis for this view by revealing that distinct L5 IT cell types participate in different interlaminar microcircuits linking layers 2/3 and 5.

Specifically, layer 2 IT neurons preferentially target L5a IT neurons, whereas layer 3 IT neurons preferentially innervate L5b-2 neurons, a pattern that cannot be explained by laminar overlap alone^57^ and is reinforced by synapse-shuffling analyses. Reciprocally, L5 IT subtypes exhibit biased connectivity back to supragranular populations, refining parallel channels across cortical depth. L5a neurons, selectively coupled to layer 2 circuits and projecting to superficial layers, are well positioned to support top-down processing, whereas L5b-2 neurons form recurrent networks with layer 3 circuits consistent with bottom-up processing. L5b-1 neurons exhibit more integrative connectivity, bridging both pathways.

This organization indicates that supragranular inputs are not collapsed into a single representation within layer 5 IT neurons. Instead, L2/3 and L5 IT neurons form cell-type-specific recurrent channels that preserve information stream identity while enabling selective transformation through feedback-like interactions. Channel convergence occurs most prominently at the L5 ET neurons (but also at the L5b-1), which receive convergent input from multiple IT pathways. Importantly, this integration remains structured rather than uniform, as inputs from layer 2 and layer 3 target distinct dendritic compartments of ET neurons, and different L5 IT types engage ET cells in distinct ways.

Within predictive processing frameworks, this circuit architecture provides a plausible substrate for parallel handling of sensory evidence and contextual predictions. L2-biased circuits carrying contextual information and L3-biased circuits carrying sensory evidence can both engage L5 IT neurons, which in turn provide channel-specific feedback to modulate supragranular processing without mutual interference. In contrast, ET neurons appear to function as output integrators that consolidate information for downstream targets. Thus, the sequential engagement of IT- and ET-dominated circuits may implement a progression from channel-specific cortical processing to action- or behavior-relevant output.

Consistent with this interpretation, L5 IT cell types with morphologies corresponding to those identified here exhibit distinct long-range projection patterns^24,52^ (Fig. 6d, Extended Data Fig. 5), with L5b-2-like neurons largely restricting their projections to higher visual areas whereas L5a-and L5b-1–like neurons projecting more broadly to higher-order cortical and subcortical regions. Furthermore, antipsychotic drugs reduce activity correlations to different extents across molecularly distinct L5 IT subpopulations throughout dorsal cortical areas, including VISp, indicating cell-type-specific modulation across the cortex^58,59^. These observations suggest that L5 IT channel-specific organization extends beyond local circuitry to brain-wide networks. Together, these findings position L5 IT neurons not as generic integrators, but as channel-specific relays that preserve and transform parallel streams of cortical information before their convergence at cortical output stages.

### Inhibition as a channel-preserving operator in cortical circuits

Inhibition plays a central role in shaping how these information channels interact. Rather than acting as a global normalizing mechanism or mediating competition across top-down and bottom-up channels, our results support a channel-preserving manner of inhibition across supragranular and infragranular circuits. Specifically, strong recurrent networks are formed in layer 2/3 and layer 5 between inhibitory neurons and the same excitatory cell type, or cell, that engages them. Moreover, L2/3 and L5 IT neurons frequently target the same postsynaptic cells through both direct synaptic connections and disynaptic connections mediated by inhibitory neurons. This connectivity pattern, commonly described as “feedforward inhibition”, enforces brief integration windows and enables precise coincidence detection^49^. With inhibition custom-matched to excitatory identity, each excitatory pathway remains independently regulated, preserving its directionality and input–output relationships while minimizing cross-channel interference. In addition, reciprocal inhibitory loops provide rapid, channel-specific feedback that supports timing control and gain modulation while stabilizing recurrent excitation. By preferentially recruiting matched inhibitory partners, each excitatory population receives immediate and proportional feedback, enabling stable amplification without runaway activity. Notably, similar circuit configurations mediated by distinct inhibitory neuron types can support different functions. For example, parvalbumin (PV)- and somatostatin (SST)-expressing interneurons (enriched in our PTC and DTC groups, respectively) differ in physiology and subcellular targeting^25,60–62^, and contribute differently to L2/3 IT receptive fields and orientation selectivity^6,63,64^. Both PTCs and DTCs are strongly recruited across the circuits examined in this study, likely enabling diverse and fine-tuned regulation of neuronal activity within individual channels.

This channel-specific organization provides insight into predictive processing frameworks, in which top-down predictions and bottom-up errors are processed in parallel, and inhibition must prevent mutual interference while still allowing interaction. Channel-matched inhibition provides exactly this capability. It enables parallel streams to remain segregated during early processing, while allowing selective coordination through higher-order loops, such as those mediated by L5 IT neurons. It also suggests that selective perturbation of specific inhibitory motifs could disrupt defined computations—such as contextual modulation or sensory integration—without globally destabilizing the network.

A requirement of some predictive coding models is the presence of cross-channel inhibition, whereby activity in one processing stream can selectively suppress or modulate another^45^. Although cross-channel inhibition is not prominent at the population level in our study, analysis at single-cell resolution reveals potential substrates. For example, a subset of layer 5 DTC neurons within inhibitory motif 9b selectively target either L2 or L3 IT neurons, indicating a possible mechanism for fine-scale, cross-channel interactions. While the number of neurons proofread in the dataset limits a systematic assessment of their recruitment, more complete datasets will be needed to determine whether these neurons support dynamic, input-dependent coordination between parallel cortical channels.

### A canonical grammar

The concept of a canonical cortical microcircuit has long occupied an uneasy position in systems neuroscience. On one hand, cortical areas display strikingly conserved architectural motifs, suggesting the existence of shared organizing principles. On the other hand, the diversity of cell types, connectivity patterns, and functional specializations across cortical areas has made it easy to portray the canonical microcircuit as either too abstract to be informative or too rigid to accommodate biological variability. In this study, we engage this tension directly by taking the canonical microcircuit proposed by Douglas and Martin (1989)^1^ as a starting point and asking how its core principles are instantiated when examined at synaptic resolution.

The original canonical microcircuit identified a set of recurring interaction motifs, including recurrent excitation within cortical layers, structured recurrent exchanges across layers, and inhibitory networks that stabilize and shape activity. Our findings are broadly consistent with this architecture, but refine it in a critical way. Rather than collapsing these motifs into a single recurrent pool, we find that the cortex implements them in spatially overlapping, cell-type-specific channels. Each channel preserves the core features of the canonical circuit—recurrent excitation, interlaminar coupling, and inhibitory control—while exhibiting distinct connectivity biases that suggest the routing of different streams of information.

Specifically, we identify two overlapping supragranular channels biased toward L2 and L3 IT neurons, respectively. Both channels exhibit robust recurrent excitation and inhibition, and both engage interlaminar loops involving L5 IT neurons in a cell-type-specific manner. In this architecture, information flow is routed, gated, and refined, not simply pooled onto L5 output pathways.

As emphasized in earlier formulations of the canonical microcircuit, cortical circuits are not static entities but transient configurations formed by subsets of neurons whose activity exceeds an inhibitory threshold. Our findings clarify how wiring rules constrain such dynamics. Depth-dependent recurrence restricts which neurons preferentially amplify one another, cell-type-specific reciprocity determines which recurrent loops are reinforced, and matched inhibition sets the conditions under which activity sustains itself or dissolves. Together, these constraints allow multiple functional circuits to coexist within the same cortical volume without anatomical segregation. In this sense, the canonical microcircuit is better understood as defining a grammar of connectivity—a set of rules governing recurrence, inhibition, reciprocity, and subcellular targeting—from which diverse circuit implementations and dynamic neuronal assemblies can be generated within the same cortical space.

## Methods

### MICrONS dataset

This study used a large-scale serial-section transmission electron microscopy (ssTEM) volume of mouse visual cortex, generated as part of the MICrONS project^32^. Detailed descriptions of tissue preparation, image acquisition, and data processing are provided in the MICrONS associated publication package^32^. Briefly, prior to EM processing, the animal (postnatal day 78) underwent in vivo two-photon calcium imaging to record visually evoked activity from excitatory neurons across cortical layers within a region spanning the junction of primary visual cortex (VISp) and three higher visual areas — lateromedial (VISlm), rostrolateral (VISrl), and anterolateral (VISal). Following functional imaging, the same region was prepared for ssTEM, enabling dense reconstruction of neuronal morphology and synaptic connectivity throughout the imaged volume.

All analyses in this study were performed on a subvolume of this image dataset, which contains 65% of the sections with dimensions of 0.52 × 1.1 × 0.82 mm (anteroposterior × mediolateral × depth). The data resources are publicly available at https://www.microns-explorer.org/.

### Visual cortical area assignment for individual cells

As part of the *in vivo* functional data collection, area borders between primary visual cortex (VISp) and higher order visual areas (HVA) were determined by retinotopic sign mapping^32^. Further work coregistered the MICrONS functional data and electron microscopy image stacks by applying a spline transform between the data stacks, anchored by vasculature landmarks. Manually-validated coregistration of n=15,352 individual EM reconstructed neurons matched to their two-photon counterpart provides a high-confidence linkage between data stacks. The coregistration information is available from the Connectome Annotation Versioning Engine (CAVE)^65^ table: coregistration_manual_v4.

Area assignment for all EM reconstructed cells with a nucleus (n=144,120) is computationally inferred from this coregistered subset. First, all nucleus positions were transformed into flattened coordinates, removing a 5-degree tilt of the pial surface (package: standard_transform https://github.com/CAVEconnectome/standard_transform). Then all positions were projected in a top-down view (XZ view in the EM space, XY view in two-photon space). The coregistered positions were labeled with their visual area assignments (V1 – primary visual area; RL – rostrolateral visual area; AL – anterolateral visual area; LM - lateromedial visual area). A support vector machine (SVM) trained on the coregistered positions then drew the area boundaries on the two-dimensional plane. All nuclei in the dataset were assigned an inferred brain area based on their XZ position relative to the area boundaries. The distance in µm to the nearest boundary is provided as a confidence measure. Area assignment and distance are available from the CAVE table: nucleus_functional_area_assignment.

### Cell type classification and correction

A hierarchical classifier based on perisomatic features was trained to predict cell types across the dataset^36^. The model was trained using cell type labels derived from data-driven clustering of column neurons as described in a previous publication^3^, and a total of 71,987 neurons in the dataset have cell types identified. Excitatory neurons were classified into 17 M-types with different morphological and synaptic features. Inhibitory neurons were grouped into four subclasses distinguished by their synaptic features, subcellular targeting patterns, and preferred postsynaptic cell types. The classifier did not assign any cell to the L6wm type, as many neurons in the white matter failed quality control for feature extraction, likely due to alignment quality in the white matter. Cell types of all neurons are available from CAVE table: aibs_metamodel_mtypes_v661_v2.

In cases of disagreement between whole-dataset predictions and column-based annotations, the column-derived cell type was used. In addition, the whole-dataset predictions were compared against manually labeled ground truth^36^. Cells without consensus in excitatory/inhibitory classification were reviewed by human experts to determine cell types based on the morphological and synaptic properties of dendritic arbors.

### Thalamocortical axon identification

Putative axons from the dorsal lateral geniculate nucleus (dLGN) were identified based on gross and fine morphological features consistent with previous reports^66–68^. These axons typically exhibit thick, myelinated shafts that enter the imaging volume via white matter bundles before ascending into gray matter, where they form a single, highly branched, and compact arbor spanning layers 2/3 and 4. Each axon forms numerous large boutons, making asymmetric (putative excitatory) synapses. Their strong synaptic connectivity in layer 4 further supports their identification as thalamocortical inputs.

### Manual proofreading strategies and status

Proofreading of neuronal reconstructions was done by trained proofreaders from the Allen Institute and Six Eleven Global Services, supported by the Virtual Observatory of the Cortex (VOrtex) program (https://www.microns-explorer.org/vortex). The proofreading process was performed as previously described ^3,32^ using the CAVE infrastructure ^65^ and a modified version of Neuroglancer with annotation capabilities, which enabled manual split and merge edits of segmented neurons. Proofreaders were aided by on-demand highlighting of branch points and terminal tips within user-defined regions based on rapid skeletonization (https://github.com/AllenInstitute/Guidebook, and a newer version https://github.com/CAVEconnectome/Tourguide), allowing efficient identification of potential false merges, extension opportunities, and systematic tracking of reviewed arbor regions.

The proofreading strategies and status of individual cells are documented at https://tutorial.microns-explorer.org/proofreading.html. For dendrites of all the column cells and nearby cells analyzed for output connectivity, all branch points were examined for correctness, and terminal tips were assessed for possible extension. Detached spine heads were not comprehensively proofread, and previous estimates place the rate of detachment at roughly 10%. For axons, the false mergers were also removed by systematically inspecting all the branch points, but the strategies for axonal extension varied by neuron type. The status for cells included for output connectivity analysis is as follows:

1. For all the comprehensively proofread L2/3 IT (n=45 cells), L4 IT (n=115 cells), L5 IT (n=55), and L5 ET (n=7)^31^ neurons (referred to as ‘proofread’ in this study), every terminal tip of axon was manually inspected and extended as far as possible, until reaching either a true biological end or an incomplete end the expert annotators could not confidently trace beyond. Incomplete ends occurred when axons reached the dataset border or were truncated by imaging artifacts. The analyses mainly focused on these neurons.
2. In addition to those comprehensively extended neurons, L2/3 IT (n=322) and L4 IT (n=189) neurons within the VISp column were reconstructed to varying levels of completion, at least to the point where the primary axon branch and local collaterals crossed the boundary of the 100 μm-wide cortical column.
3. A total of 413 proofread inhibitory neurons within and near the column received substantial axonal extension as described previously^3^.

Several dataset-specific limitations should be noted. Incomplete segmentation in the upper ∼10 µm of layer 1 truncated portions of apical tufts and reduced reconstruction quality for layer 1 interneurons. Consequently, reconstructions of neurons with extensive apical dendritic or axonal arbors may underestimate inputs and outputs in this region.

### Connectivity analysis

To quantify the synaptic output connectivity of individual cells, we included only synapses onto segments associated with a single nucleus and assigned a defined cell type. The connectivity was represented as synapse counts or as proportions of total output synapses, across cell types or along spatial axes. To ensure reliable disynaptic connectivity measurement, analyses were restricted to inhibitory neurons with well-reconstructed axonal arbors. Weighted synapse numbers were calculated by multiplying the synapse number of the disynaptic connection by the synapse number of the monosynaptic connection. Although most of the postsynaptic targets were not proofread, estimates on the basis of proofread neurons indicate that 99% of non-proofread input synapses are accurate ^36^.

The column and halo inhibitory neurons together provided a representative sample of local inhibitory connectivity across layers, because (1) they encompassed all VISp inhibitory neuron types except Chandelier cells^3^; (2) the proofread IT neurons, located within or near the column and halo, abundantly targeted inhibitory neurons within 50µm lateral distance; (3) although the halo did not extend into L6, L2/3 IT neurons minimally targeted L6 neurons (Extended Data Figs 2b,4e).

### Inhibitory motif clustering

Proofread inhibitory neurons, except ITC neurons, were grouped into inhibitory motifs based on their synaptic output patterns within VISp, following a strategy similar to a previous study on column interneurons^3^. For each neuron, the number of output synapses onto each excitatory and inhibitory cell type was quantified and normalized. These normalized connectivity profiles were then clustered using k-means (k=19), repeated 1,000 times. A coclustering matrix was constructed, where each entry represents how often a pair of neurons was assigned to the same cluster across runs. Motif groups were then obtained by applying agglomerative clustering with complete linkage to the coclustering matrix. The number of clusters was determined by scanning solutions from 10 to 25 clusters and selecting 15 based on the silhouette score and Davies–Bouldin score, while also maintaining consistency with previously defined column inhibitory motif groups ^3^.

The resulting clusters largely recapitulated previously defined motif groups, with several notable differences: (1) the original motif 9, which selectively inhibited L5 ET neurons, was subdivided into two groups - 9a (PTC-predominant and strongly innervating inhibitory neurons) and 9b (DTC-predominant and sparsely innervating inhibitory neurons); (2) several L5 and L6 targeting motifs (8,10, 11,12, 15-18) were consolidated into four broader groups (motifs 8/10, 11/12, 15/16, and 17/18). These differences likely reflect methodological changes in the current analysis, which incorporates connectivity across the entire VISp rather than the column only, and includes inhibitory neuron targeting patterns in the clustering, whereas the previous study considered only connectivity with excitatory neurons.

### Spatial analysis

All spatial measurements were computed after a rigid rotation of 5° to flatten the pial surface and translation to zero the pial surface on the *y* axis. Cortical depth was defined by the y position of each object (soma or synapse). Radial distance was calculated as the distance between objects in the x-z plane.

### Synapse shuffling test

For each neuron, a shuffle pool was generated for every true output synapse. Specifically, a 5×5x5 um bounding box was centered on individual true output synapses to query all the synapses inside this box, together with their pre- and postsynaptic segment identities. Only synapses whose postsynaptic segments were associated with a single nucleus were included in the shuffle pool.

Shuffling was performed independently for each true output synapse by randomly sampling from its corresponding shuffle pool 1,000 times. To preserve the spatial distribution and proportion of synapses targeting excitatory and inhibitory neurons for each starting neuron, shuffle pools were separated by postsynaptic cell class (excitatory vs. inhibitory), and sampling was constrained to match the original E/I identity of the target of each synapse. All sample synapses across 1,000 iterations were aggregated and grouped by predicted cell types to calculate the average shuffled output connectivity, which was then compared to the true connectivity for each neuron.

### Feature extraction and L5 IT cell subtyping

Layer 5 IT cells were identified on the basis of clustering of a collection of dendritic features. The feature extraction pipeline was derived from the one described in ^3^ which included both skeleton features like path length and synaptic features such as size distributions and synapse depth profile envelopes, with additional perisomatic features (^3,36^ and spine features ^69^ included. UMAP dimensionality reduction and Leiden clustering on a nearest neighbors subgraph identified a group corresponding in morphology to IT cells in layers 2–5, and subsequent clustering of this group into three subgroups identified a cluster of cells centered within layer 5 that we labeled as Layer 5 IT cells. The feature depicted in Figure 4b represents a dimensionality reduction on the same collection of features, normalized only within Layer 5 IT cells. The normalization was implemented using a robust scaling process, in which each feature was centered by its median and scaled by its interquartile range to reduce sensitivity to outliers. After scaling, the extreme high and low 0.5 percentiles were clipped. The same set of L5 IT features was used to perform Linear discriminant analysis (LDA) in Figure 4f, using manually assigned labels of the three L5 IT cell types to separate the larger L5 IT population.

### Statistical analysis

Distributions of each parameter were assessed for normality using the D’Agostino–Pearson omnibus test, and appropriate parametric or nonparametric analyses were applied accordingly. Comparisons between groups were performed using unpaired, two-tailed t-tests (parametric) or Mann–Whitney tests (nonparametric). Statistical details, including the definition and value of *n*, summary statistics (mean ± SE, unless otherwise stated), and measures of significance, are provided in the text or figure legends. Statistical significance was defined as *P* < 0.05.

## Acknowledgments

The authors thank everyone that generated and maintained the MICrONs dataset; Melissa Lerch, David Vumbaco and the Program Management team at the Allen Institute for Brain Science for their guidance for project strategy and operations; H. Zeng and Clay R. Reid for their support and leadership; Agnes Bodor for discussions on cell types; Georg Keller and Anne Vasilevskaya for valuable discussions on the theoretical and computational implications of the data; we thank Shane Wilson for comments on the manuscript. Aspects of this project originated from interactive discussions and hands-on working groups at the CapoCaccia Neuromorphic Intelligence Workshop (https://capocaccia.cc), which the authors gratefully acknowledge.

The work was supported by the NIH BRAIN CONNECTS grant UM1NS132253, the NIH BICAN grant RF1MH128840 and the NIH VOrtex grant: U24NS120053. This research was supported by the Allen Institute, founded by Jody Allen – chair and co-founder of Allen Family Philanthropies, and the late Paul G. Allen – investor, philanthropist, and co-founder of Microsoft. We gratefully acknowledge their vision and generosity, which make this work possible.

## Author contributions

C.Z. contributed to conceptualization, data curation, proofreader management, data analysis, writing of the original draft, and reviewing and editing of the manuscript. C.M.S.M. contributed to data curation, data analysis, writing of the original draft, development of data infrastructure and analytical tools, and reviewing and editing of the manuscript. B.P.D. contributed to proofreader management, development of data infrastructure and analytical tools, and reviewing and editing of the manuscript. R.S. contributed to data curation. E.N. contributed to data curation. E.J. contributed to development of analytical tools. B.D.P. contributed to development of data infrastructure and analytical tools, and reviewing and editing of the manuscript. F.C.C. contributed to development of data infrastructure and analytical tools, reviewing and editing of the manuscript, and funding acquisition. N.M.d.C. contributed to conceptualization, writing of the original draft, reviewing and editing of the manuscript, and funding acquisition.

## References

1. Douglas, R. J., Martin, K. A. C. & Whitteridge, D. A canonical microcircuit for neocortex. Neural Comput. 1, 480–488 (1989).

2. Binzegger, T., Douglas, R. J. & Martin, K. A. C. A quantitative map of the circuit of cat primary visual cortex. J. Neurosci. 24, 8441–8453 (2004).

3. Schneider-Mizell, C. M. et al. Inhibitory specificity from a connectomic census of mouse visual cortex. Nature 640, 448–458 (2025).

4. Sievers, M. et al. Connectomic reconstruction of a cortical column. bioRxiv 2024.03.22.586254 (2024) doi:10.1101/2024.03.22.586254.

5. Narayanan, R. T. et al. Beyond columnar organization: Cell type- and target layer-specific principles of horizontal axon projection patterns in rat vibrissal cortex. Cereb. Cortex 25, 4450–4468 (2015).

6. Niell, C. M. & Scanziani, M. How Cortical Circuits Implement Cortical Computations: Mouse Visual Cortex as a Model. Annual Review of Neuroscience vol. 44 517–546 Preprint at 10.1146/annurev-neuro-102320-085825 (7 2021).

7. Weiler, S. et al. Functional and structural features of L2/3 pyramidal cells continuously covary with pial depth in mouse visual cortex. Cereb. Cortex 33, 3715–3733 (4 2023).

8. Tasic, B. et al. Shared and distinct transcriptomic cell types across neocortical areas. Nature 563, 72–78 (11 2018).

9. Cheng, S. et al. Vision-dependent specification of cell types and function in the developing cortex. Cell 185, 311–327.e24 (1 2022).

10. O’Toole, S. M., Oyibo, H. K. & Keller, G. B. Molecularly targetable cell types in mouse visual cortex have distinguishable prediction error responses. Neuron 111, 2918–2928.e8 (9 2023).

11. Weis, M. A. et al. An unsupervised map of excitatory neuron dendritic morphology in the mouse visual cortex. Nat. Commun. 16, (12 2025).

12. Young, H., Belbut, B., Baeta, M. & Petreanu, L. Laminar-specific cortico-cortical loops in mouse visual cortex. Elife 10, 1–25 (2 2021).

13. Morgenstern, N. A., Bourg, J. & Petreanu, L. Multilaminar networks of cortical neurons integrate common inputs from sensory thalamus. Nat. Neurosci. 19, 1034–1040 (2016).

14. Jordan, R. & Keller, G. B. Opposing influence of top-down and bottom-up input on excitatory layer 2/3 neurons in mouse primary visual cortex. Neuron 108, 1194–1206.e5 (2020).

15. Vasilevskaya, A. & Keller, G. B. A functional influence based circuit motif that constrains the set of plausible algorithms of cortical function. Preprint at 10.64898/2026.01.29.702557 (1 2026).

16. Brandalise, F. et al. Thalamocortical feedback selectively controls pyramidal neuron excitability. Nat. Commun. 16, (12 2025).

17. Bureau, I., Paul, F. S. V. & Svoboda, K. Interdigitated paralemniscal and lemniscal pathways in the mouse barrel cortex. PLoS Biol. 4, 2361–2371 (2006).

18. Oberlaender, M. et al. Three-dimensional axon morphologies of individual layer 5 neurons indicate cell type-specific intracortical pathways for whisker motion and touch. Proc. Natl. Acad. Sci. U. S. A. 108, 4188–4193 (2011).

19. Petreanu, L., Mao, T., Sternson, S. M. & Svoboda, K. The subcellular organization of neocortical excitatory connections. Nature 457, 1142–1145 (2 2009).

20. Xu, X. et al. Primary visual cortex shows laminar-specific and balanced circuit organization of excitatory and inhibitory synaptic connectivity. J. Physiol. 594, 1891–1910 (4 2016).

21. Hage, T. A. et al. Synaptic connectivity to L2/3 of primary visual cortex measured by two-photon optogenetic stimulation. Elife 11, (1 2022).

22. Patiño, M., Rossa, M. A., Lagos, W. N., Patne, N. S. & Callaway, E. M. Transcriptomic cell-type specificity of local cortical circuits. Neuron (2024) doi:10.1016/j.neuron.2024.09.003.

23. Yao, S. et al. A whole-brain monosynaptic input connectome to neuron classes in mouse visual cortex. Nat. Neurosci. 26, 350–364 (12 2022).

24. Sorensen, S. A. et al. Connecting single-cell transcriptomes to projectomes in mouse visual cortex. Nature (2026) In Press. bioRxiv (2023) doi:10.1101/2023.11.25.568393

25. Gouwens, N. W. et al. Integrated Morphoelectric and Transcriptomic Classification of Cortical GABAergic Cells. Cell 183, 935–953.e19 (11 2020).

26. Petilla Interneuron Nomenclature Group et al. Petilla terminology: nomenclature of features of GABAergic interneurons of the cerebral cortex. Nat. Rev. Neurosci. 9, 557–568 (2008).

27. Kubota, Y. Untangling GABAergic wiring in the cortical microcircuit. Curr. Opin. Neurobiol. 26, 7–14 (2014).

28. Isaacson, J. S. & Scanziani, M. How inhibition shapes cortical activity. Neuron 72, 231–243 (2011).

29. Lee, W.-C. A. et al. Anatomy and function of an excitatory network in the visual cortex. Nature 532, 370–374 (2016).

30. Seeman, S. C. et al. Sparse recurrent excitatory connectivity in the microcircuit of the adult mouse and human cortex. Elife 7, (2018).

31. Bodor, A. L. et al. The synaptic architecture of layer 5 thick tufted excitatory neurons in mouse visual cortex. Nat. Neurosci. 28, 1704–1715 (2025).

32. MICrONS Consortium. Functional connectomics spanning multiple areas of mouse visual cortex. Nature 640, 435–447 (2025).

33. Gamlin, C. R. et al. Connectomics of predicted Sst transcriptomic types in mouse visual cortex. Nature 640, 497–505 (2025).

34. Tasic, B. et al. Adult mouse cortical cell taxonomy revealed by single cell transcriptomics. Nat. Neurosci. 19, 335–346 (2016).

35. Kim, E. J., Juavinett, A. L., Kyubwa, E. M., Jacobs, M. W. & Callaway, E. M. Three Types of Cortical Layer 5 Neurons That Differ in Brain-wide Connectivity and Function. Neuron 88, 1253–1267 (2015).

36. Elabbady, L. et al. Perisomatic ultrastructure efficiently classifies cells in mouse cortex. Nature 640, 478–486 (2025).

37. Ko, H. et al. Functional specificity of local synaptic connections in neocortical networks. Nature 473, 87–91 (5 2011).

38. Ding, Z. et al. Functional connectomics reveals general wiring rule in mouse visual cortex. bioRxiv 2023.03.13.531369 (3 2023).

39. Campagnola, L. et al. Local connectivity and synaptic dynamics in mouse and human neocortex. Science 375, (3 2022).

40. Oldenburg, I. A. et al. The logic of recurrent circuits in the primary visual cortex. Nat. Neurosci. 27, 137–147 (1 2024).

41. Bock, D. D. et al. Network anatomy and in vivo physiology of visual cortical neurons. Nature 471, 177–182 (2011).

42. Bopp, R., da Costa, N. M., Kampa, B. M., Martin, K. A. C. & Roth, M. M. Pyramidal Cells Make Specific Connections onto Smooth (GABAergic) Neurons in Mouse Visual Cortex. PLoS Biology vol. 12 e1001932 Preprint at 10.1371/journal.pbio.1001932 (2014).

43. Znamenskiy, P. et al. Functional specificity of recurrent inhibition in visual cortex. Neuron 112, 991–1000.e8 (2024).

44. Ogando, M. B., et al. Feature-specific inhibitory connectivity augments the accuracy of cortical representations. bioRxivorg (2025) doi:10.1101/2025.08.02.668307.

45. Keller, G. B. & Mrsic-Flogel, T. D. Predictive Processing: A Canonical Cortical Computation. Neuron 100, 424–435 (2018).

46. Gilbert, C. D. & Wiesel, T. N. Morphology and intracortical projections of functionally characterised neurones in the cat visual cortex. Nature 280, 120–125 (1979).

47. Bastos, A. M. et al. Canonical microcircuits for predictive coding. Neuron 76, 695–711 (2012).

48. Moberg, S. & Takahashi, N. Neocortical layer 5 subclasses: From cellular properties to roles in behavior. Front. Synaptic Neurosci. 14, 1006773 (2022).

49. Pouille, F. & Scanziani, M. Enforcement of temporal fidelity in pyramidal cells by somatic feed-forward inhibition. Science 293, 1159–1163 (2001).

50. Zeng, H. & Sanes, J. R. Neuronal cell-type classification: challenges, opportunities and the path forward. Nat. Rev. Neurosci. 18, 530–546 (2017).

51. Scala, F. et al. Phenotypic variation of transcriptomic cell types in mouse motor cortex. Nature 598, 144–150 (2021).

52. Gao, L. et al. Integrative analysis of single-neuron projectomes links connectome, transcriptome, and function in the mouse cortex. Neuron 114, 86–104.e5 (2026).

53. Rao, R. P. & Ballard, D. H. Predictive coding in the visual cortex: a functional interpretation of some extra-classical receptive-field effects. Nat. Neurosci. 2, 79–87 (1999).

54. Aizenbud, I., et al. Neural mechanisms of predictive processing: a collaborative community experiment through the OpenScope program. arXiv [q-bio.NC] (2026) doi:10.48550/arXiv.2504.09614.

55. Larkum, M. A cellular mechanism for cortical associations: an organizing principle for the cerebral cortex. Trends Neurosci. 36, 141–151 (2013).

56. Onodera, K. & Kato, H. K. Translaminar recurrence from layer 5 suppresses superficial cortical layers. Nat. Commun. 13, (12 2022).

57. Matho, K. S. et al. Genetic dissection of the glutamatergic neuron system in cerebral cortex. Nature 598, 182–187 (10 2021).

58. Heindorf, M. & Keller, G. B. Antipsychotic drugs selectively decorrelate long-range interactions in deep cortical layers. Elife 12, RP86805 (2024).

59. Lupori, L. et al. A single dose of the antipsychotic drug clozapine has long-term behavioral and functional effects in mice. bioRxiv 2026.03.27.714783 (2026) doi:10.64898/2026.03.27.714783.

60. Tremblay, R., Lee, S. & Rudy, B. GABAergic interneurons in the neocortex: From cellular properties to circuits. Neuron 91, 260–292 (2016).

61. Kepecs, A. & Fishell, G. Interneuron cell types are fit to function. Nature 505, 318–326 (2014).

62. Morabito, A. et al. Distinct dendritic integration strategies control dynamics of inhibition in the neocortex. Neuron 113, 2962–2978.e10 (2025).

63. Nienborg, H. et al. Contrast dependence and differential contributions from somatostatin- and parvalbumin-expressing neurons to spatial integration in mouse V1. J. Neurosci. 33, 11145–11154 (2013).

64. Adesnik, H., Bruns, W., Taniguchi, H., Huang, Z. J. & Scanziani, M. A neural circuit for spatial summation in visual cortex. Nature 490, 226–231 (2012).

65. Dorkenwald, S. et al. CAVE: Connectome Annotation Versioning Engine. Nat. Methods 22, 1112–1120 (2025).

66. Antonini, A., Fagiolini, M. & Stryker, M. P. Anatomical correlates of functional plasticity in mouse visual cortex. J. Neurosci. 19, 4388–4406 (1999).

67. Peng, H. et al. Morphological diversity of single neurons in molecularly defined cell types. Nature 598, 174–181 (10 2021).

68. Bopp, R., Holler-Rickauer, S., Martin, K. A. C. & Schuhknecht, G. F. P. An ultrastructural study of the thalamic input to layer 4 of primary motor and primary somatosensory cortex in the mouse. J. Neurosci. 37, 2435–2448 (2017).

69. Pedigo, B. D. et al. A quantitative census of millions of postsynaptic structures in a large electron microscopy volume of mouse visual cortex. bioRxiv 2026.02.19.706834 (2026) doi:10.64898/2026.02.19.706834.

