## Extended Data Figures for "Cell-type-specific parallel pathways in the canonical cortical microcircuit"

### Extended Data Figure 1

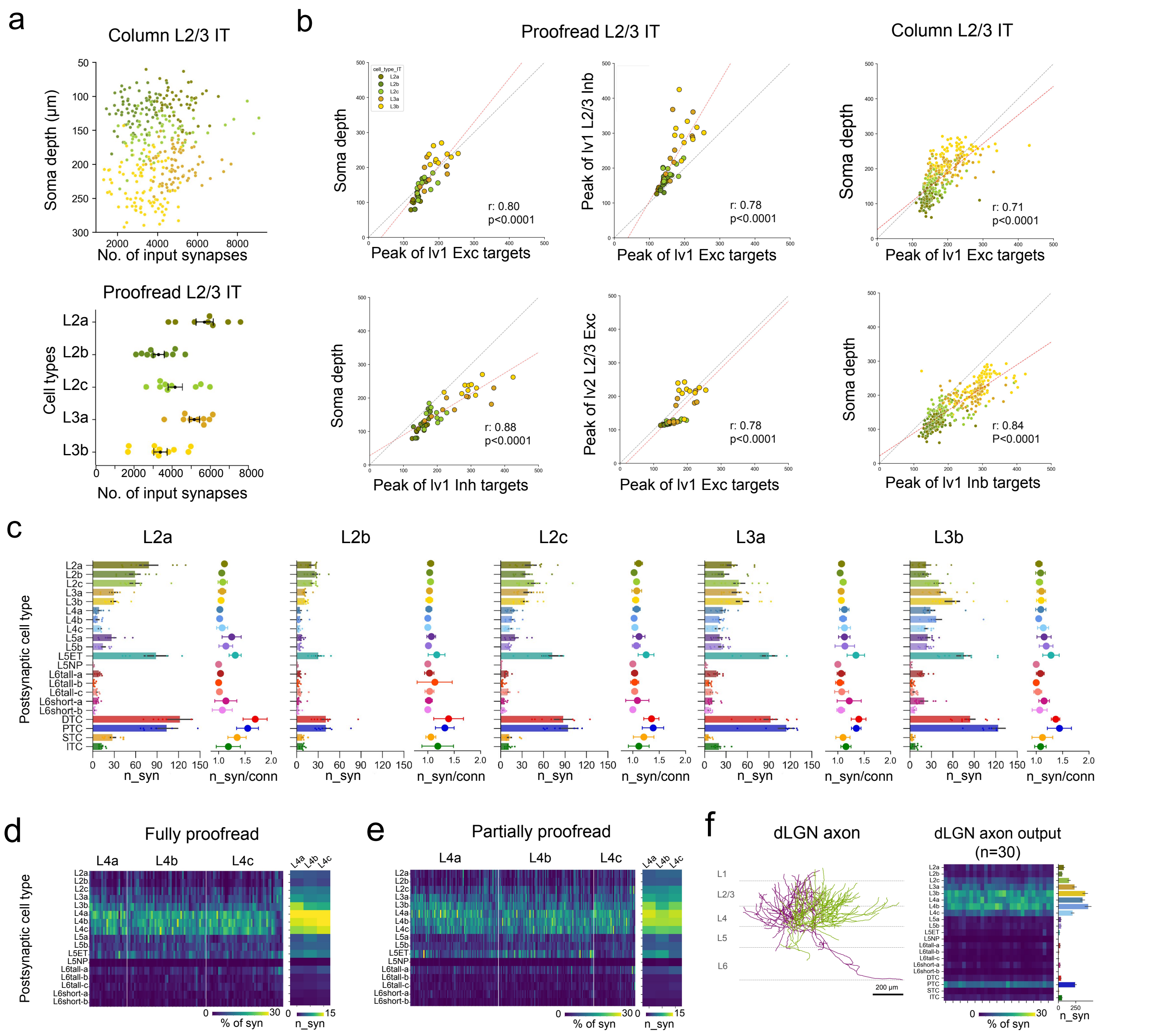


**Extended Data Figure 1**. L2/3 IT type-specific synaptic and connectivity features

**a**. Total input synapse numbers of individual column (upper panel) and proofread (lower panel) L2/3 IT neurons. **b**. Correlation analysis of presynaptic soma depth with the KDE peak of excitatory and inhibitory target soma depth distributions within layer 2/3. In each panel, the grey dashed line indicates the line of identity (x=y), and the red dashed line indicates the linear regression fit to the data points. **c**. Output connectivity of L2/3 IT neurons across cell types. Bar graphs demonstrate mean ± SE total synapse numbers made by presynaptic cells with each postsynaptic cell type. Points representing individual presynaptic cell data are overlaid on the bars. Each bar graph has a point plot to its right, demonstrating the mean ± SD number of synapses per connection made by the presynaptic L2/3 IT neuron with their postsynaptic cell types. **d**. Output connectivity of fully proofread L4 IT neurons, shown as the percentage of total output synapses across target cell types for individual neurons (left) and the average number of output synapses per target cell type (right). **e.** Same heatmap as in (d) for partially proofread L4 IT neurons. **f**. Representative skeletons of the dorsal lateral geniculate nucleus (dLGN) axons in VISp and output connectivity of proofread dLGN axons across cell types.

### Extended Data Figure 2

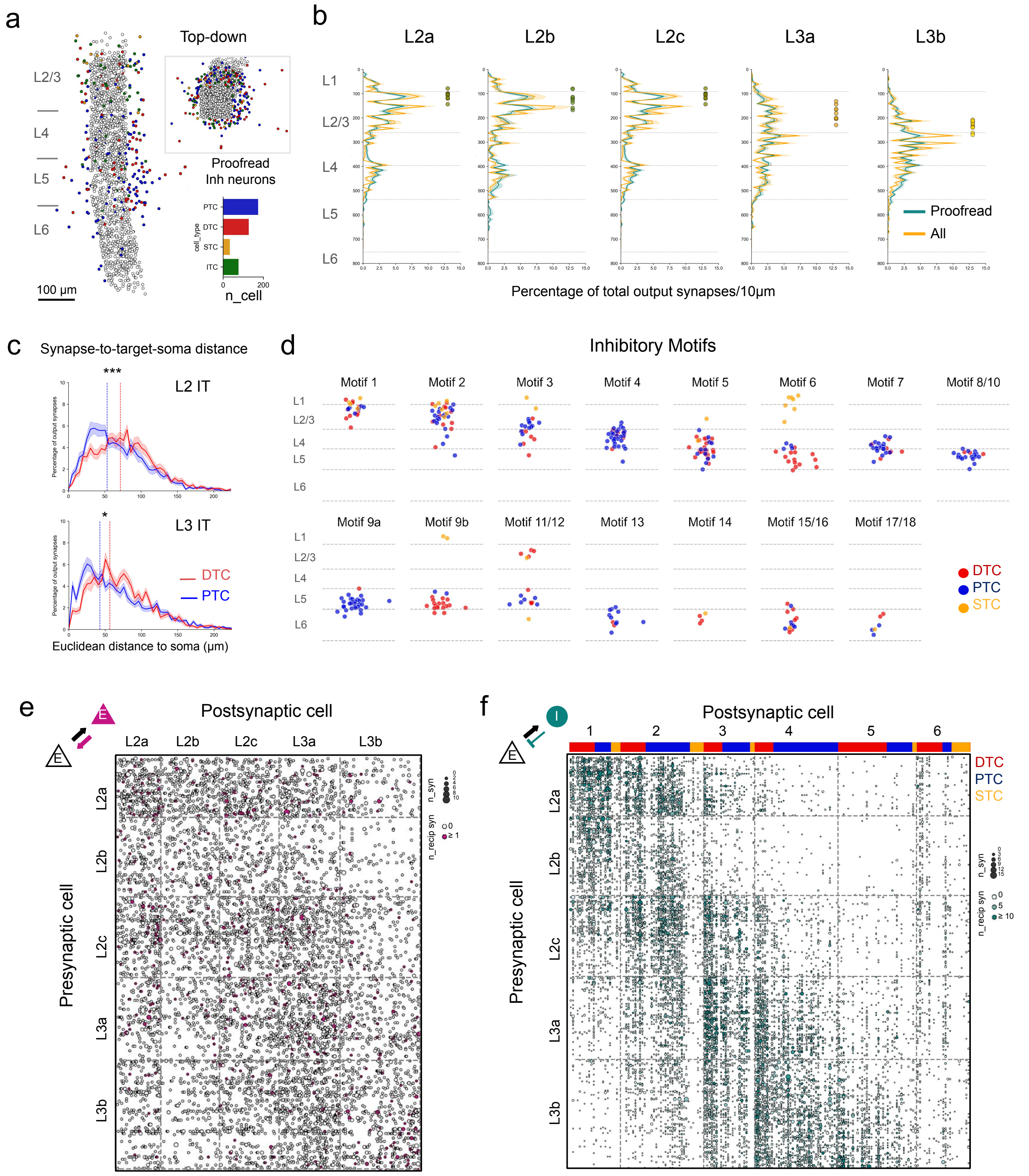

**Extended Data Figure 2**. Inhibitory motifs and reciprocal connectivity with L2/3 IT neurons

**a**. Side view (left) and top-down view (right-top) of the soma locations of the proofread column cells (open circles) and proofread inhibitory neurons surrounding the column (solid circles). The right-bottom bar graph shows the total number of proofread inhibitory neurons for each type. b. Soma depth distribution of all VISp and proofread-only inhibitory postsynaptic targets of L2/3 IT neurons. **c**. Distributions of Euclidean distances from L2 and L3 output synapses to the soma of postsynaptic PTC and DTC targets. Mean KDE peaks across individual DTC and PTC cells are indicated by color-coded dashed lines. **d**. Soma locations of inhibitory neurons from each motif across cortical layers. **e**. Connectivity matrix of individual pairs of presynaptic and postsynaptic column L2/3 IT neurons. Circle size indicates the synapse number of forward (pre-to-post) connections. Solid circles indicate the existence of reciprocal connections. **f**. Connectivity matrix of individual pairs of presynaptic column L2/3 IT neurons and postsynaptic proofread inhibitory neurons arranged by motifs and cell types (top bar). Circle size indicates the synapse number of forward (pre-to-post) connections. Circle hue indicates the synapse number of reciprocal connections. *, $p<0.05$; ***, $p<0.001$

### Extended Data Figure 3

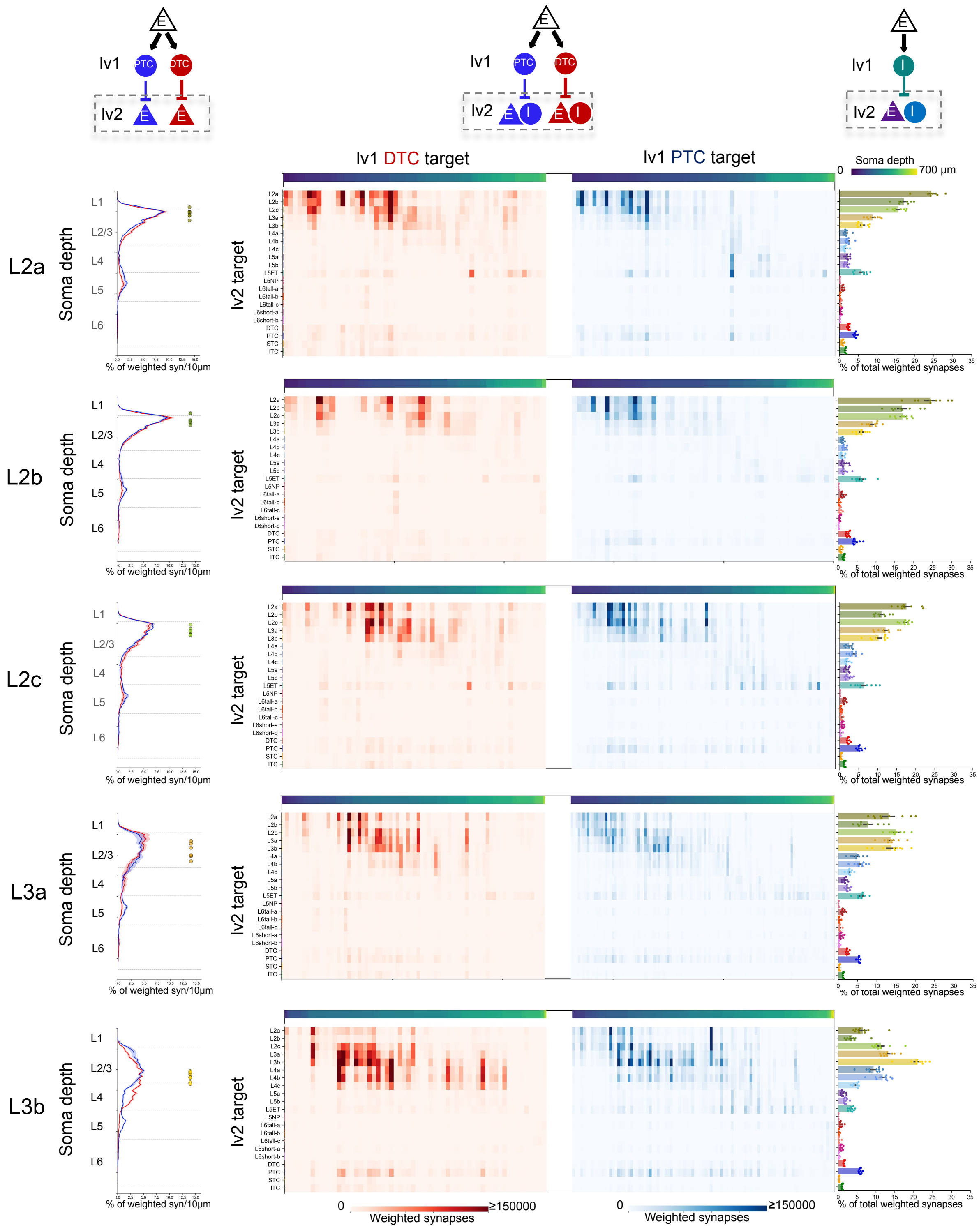


**Extended Data Figure 3.** Disynaptic connectivity of L2/3 IT neurons

Left, cortical depth distributions of disynaptic (lv2) excitatory targets postsynaptic to monosynaptic (lv1) PTC (blue) and DTC (red) targets of L2/3 IT neurons. Each line indicates the mean ± SE of the percentage of total weighted synapses at 10µm intervals across cortical depth, averaged across starting L2/3 IT neurons. Soma locations of individual starting L2/3 IT neurons are indicated by color-coded filled circles on the right of each panel; Middle, weighed output synapse numbers onto lv2 cell types of lv1 inhibitory targets postsynaptic to each L2/3 IT types. Rows are lv2 targets grouped by cell types and columns are individual lv1 inhibitory neurons arranged by soma depth (top bar); Right, percentage of total weighted synapses made between L2/3 IT types and their lv2 postsynaptic cell types. Points representing individual starting L2/3 IT neuron data are overlaid on the bars.

### Extended Data Figure 4

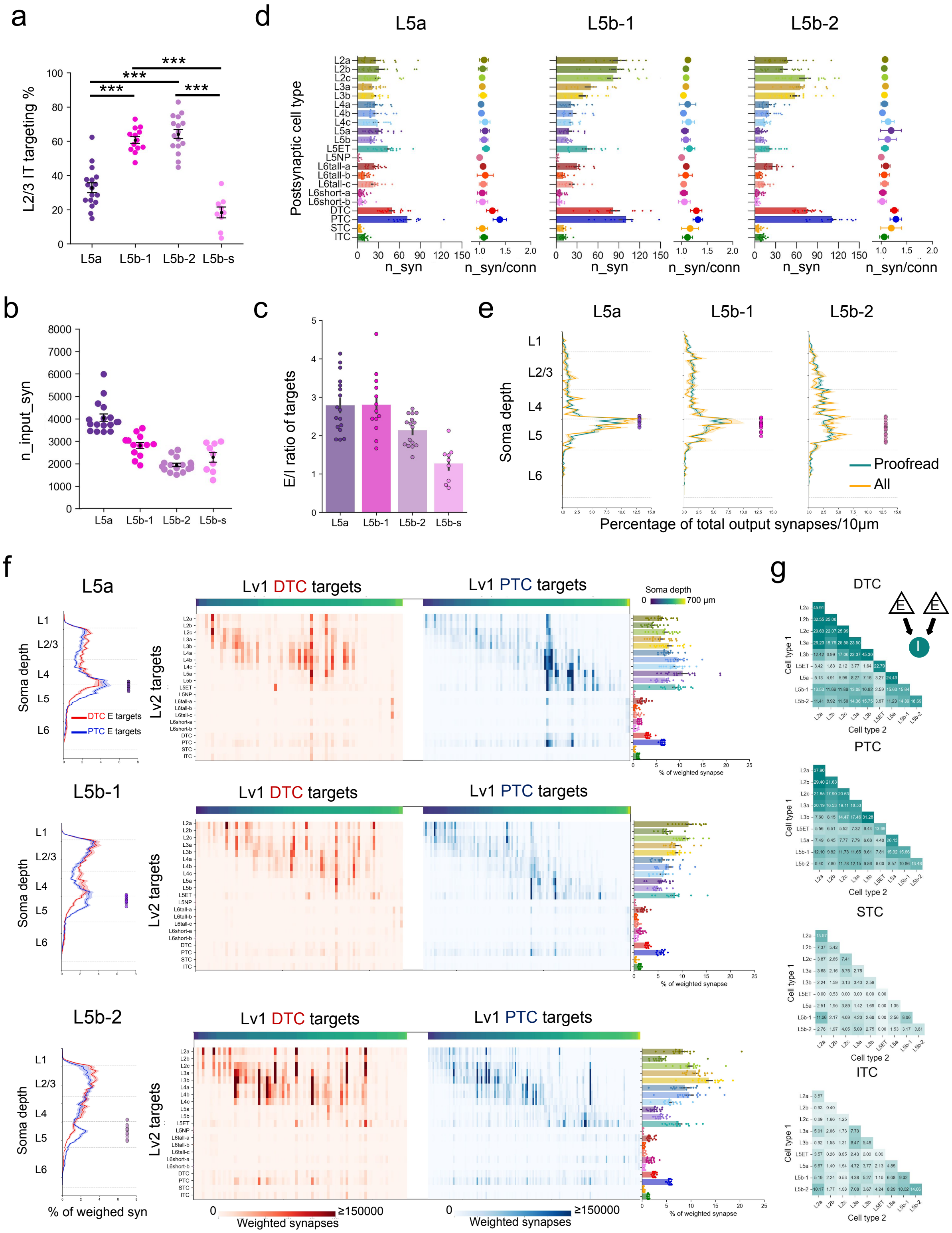

**Extended Data Figure 4.** L5 excitatory neurons have distinct synaptic features and connectivity

**a.** Percentage of L2/3 IT targeting of individual L5a and L5b IT cells. **b.** Total input synapses of individual L5 IT neurons across types. **c.** Ratio of excitatory versus inhibitory target numbers of each L5 IT type. **d.** Output connectivity of L5 IT neurons across cell types. Bar graphs demonstrate mean ± SE total synapse numbers made with each postsynaptic cell type. Points representing individual presynaptic cell data are overlaid on the bars. Each bar graph is paired with a point plot showing the mean ± SD number of synapses per connection from L5 IT neurons to each postsynaptic cell type. **e.** **f.** Left, cortical depth distributions of disynaptic (lv2) excitatory targets postsynaptic to monosynaptic (lv1) PTC (blue) and DTC (red) targets of L5 IT neurons. Middle, weighed output synapse numbers onto lv2 cell types of lv1 inhibitory targets postsynaptic to each L5 IT type. Right, percentage of total weighted synapses made between L5 IT types and their lv2 postsynaptic cell types. Points representing individual starting L5 IT neuron data are overlaid on the bars. **g.** Co-targeted rate of each inhibitory subclass within the column by pairs of excitatory neurons with the same or different types. ***, $p<0.001$

### Extended Data Figure 5

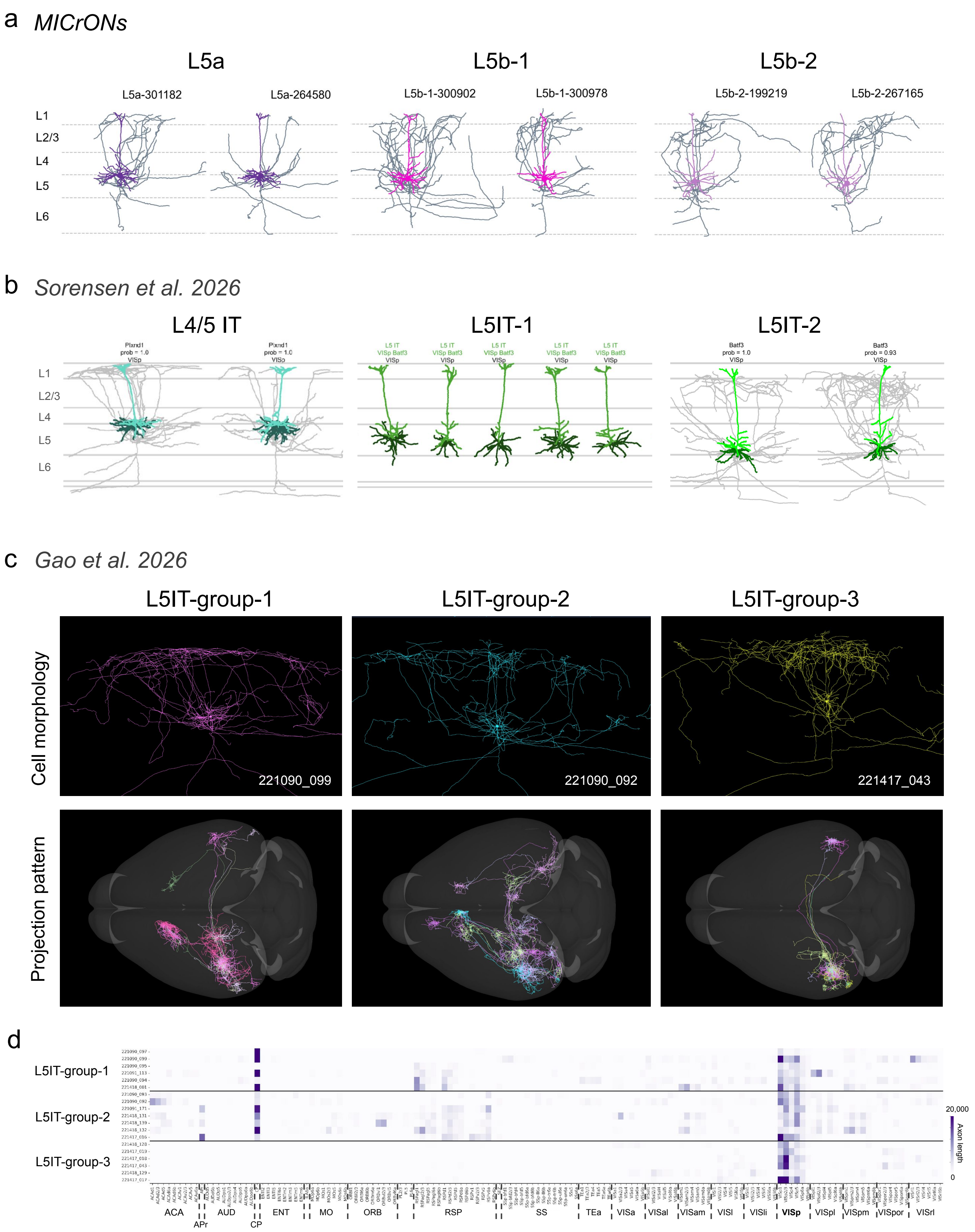

**Extended Data Figure 5.** L5 IT cell types comparison across studies

**a.** Representative morphologies of the three L5 IT types identified in this study. **b.** Adapted from Sorensen et al. 2026, showing representative morphologies of L5 IT types described in that study. **c.** Representative morphologies and long-range projection patterns of L5 IT neurons reconstructed in Gao et al., 2026, that with dendritic arbor and local axons similar to those of the L5 IT cell types identified here. Images were obtained from the publicly available data portal: https://mouse.digital-brain.cn/projectome/whole-cortex. **d.** Brain-wide projection heatmap of individual VISp L5 IT neurons shown in (c). ACA, anterior cingulate area; APr, area postriata; AUD, auditory areas; CP, caudate putamen; ENTI, entorhinal area; MO, motor area; ORB, orbital area; RSP, retrosplenial area; SS, somatosensory area; TEa, temporal association area; VISa, anterior visual area; al, anterolateral; l, lateral; li, laterointermediate; pl, posterolateral; pm, posteriomedial; por, postrhinal; rl, rostrolateral.
