## Supplementary Information for "Cell-type-specific parallel pathways in the canonical cortical microcircuit"

### Supplementary Information- Output connectivity of column excitatory neurons (CAVE v1718)

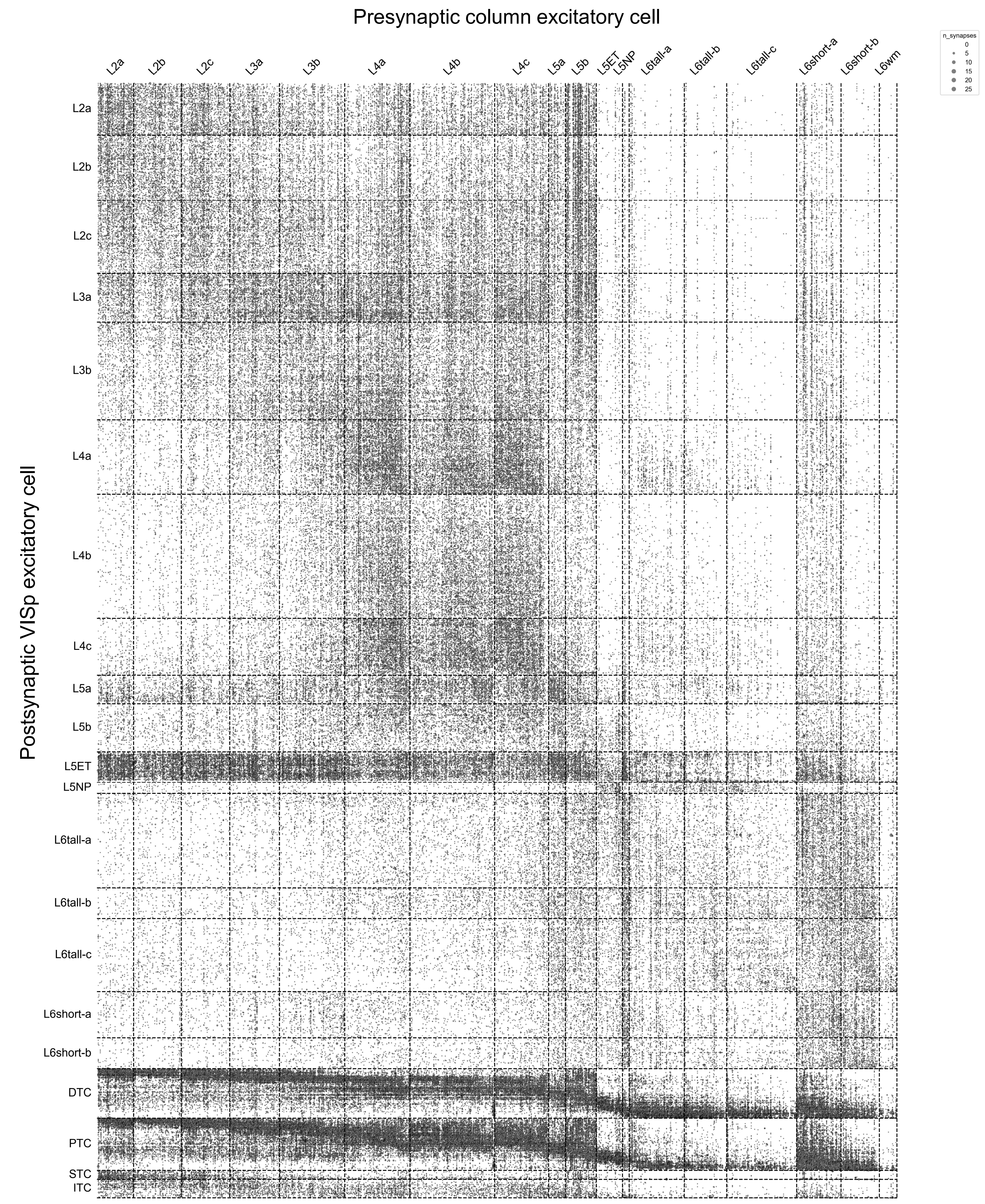

### Supplementary Information- Morphology of proofread excitatory and inhibitory neurons

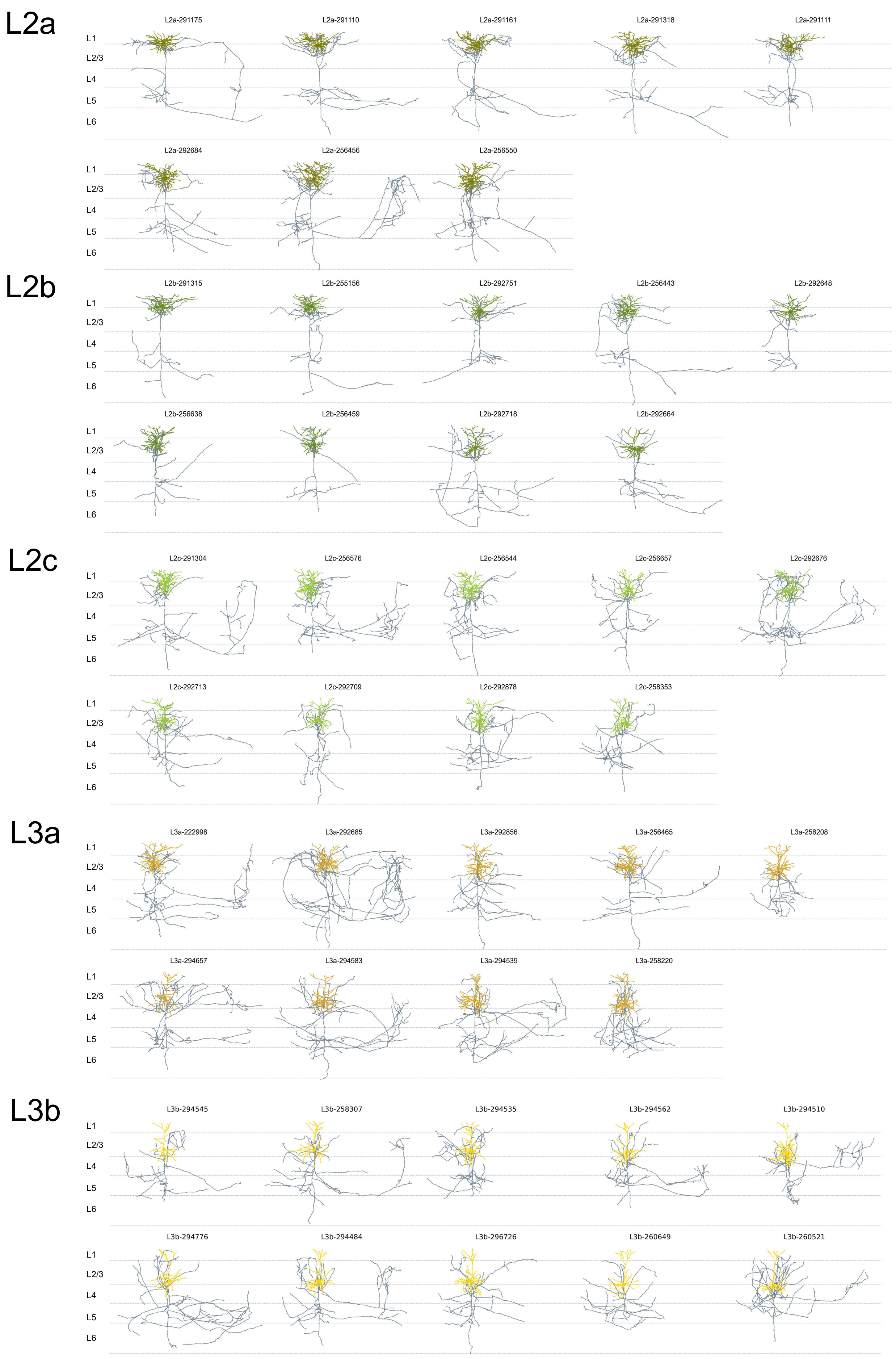

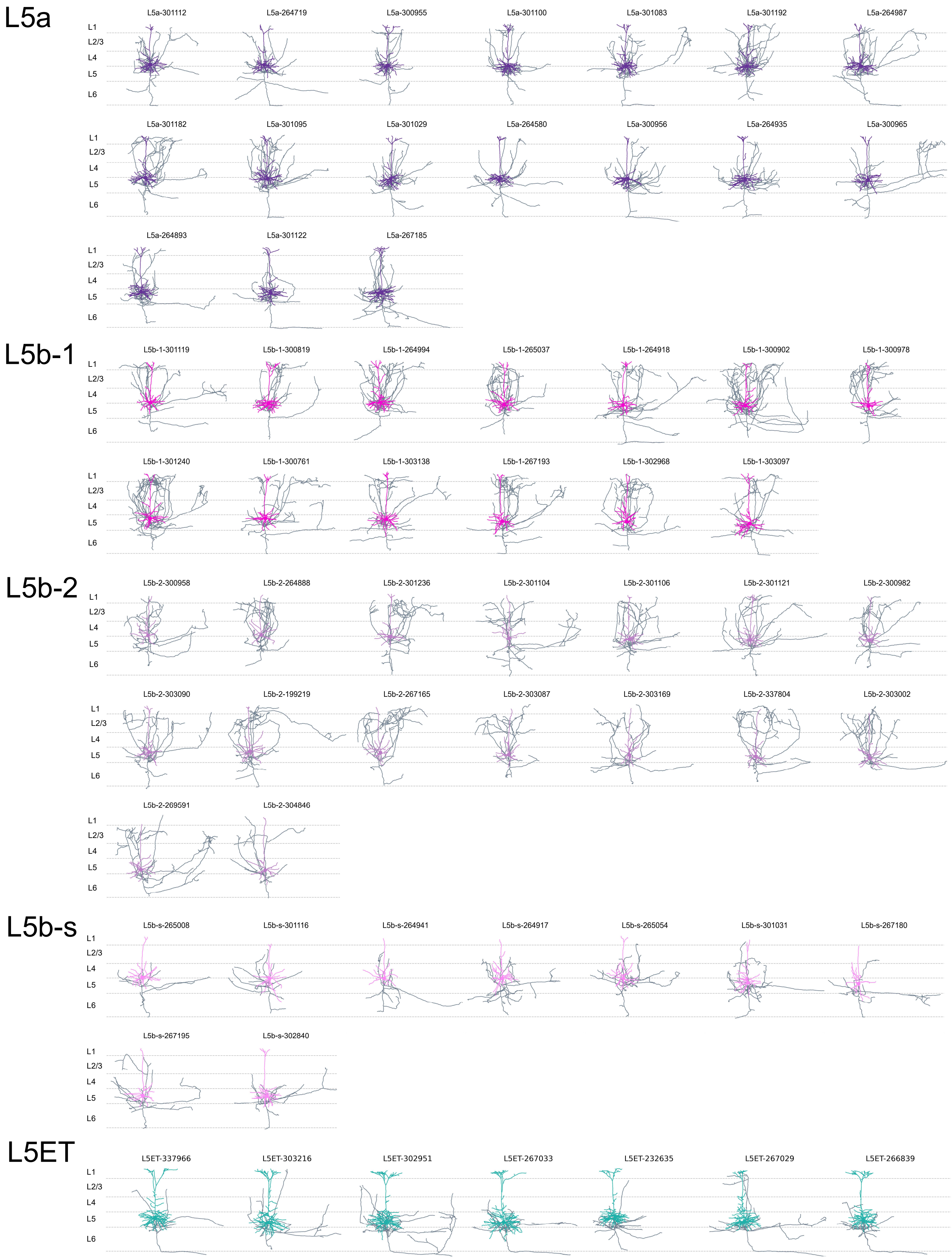
L5a
L1
L2/3
L4
L5
L6
L5a-301112
L5a-264719
L5a-300955
L5a-301100
L5a-301083
L5a-301192
L5a-264987
L5a-301182
L5a-301095
L5a-301029
L5a-264580
L5a-300956
L5a-264935
L5a-300965
L5a-264893
L5a-301122
L5a-267185
L5b-1
L5b-1-301119
L5b-1-300819
L5b-1-264994
L5b-1-265037
L5b-1-264918
L5b-1-300902
L5b-1-300978
L5b-1-301240
L5b-1-300761
L5b-1-303138
L5b-1-267193
L5b-1-302968
L5b-1-303097
L5b-2
L5b-2-300958
L5b-2-264888
L5b-2-301236
L5b-2-301104
L5b-2-301106
L5b-2-301121
L5b-2-300982
L5b-2-303090
L5b-2-199219
L5b-2-267165
L5b-2-303087
L5b-2-303169
L5b-2-337804
L5b-2-303002
L5b-2-269591
L5b-2-304846
L5b-s
L5b-s-265008
L5b-s-301116
L5b-s-264941
L5b-s-264917
L5b-s-265054
L5b-s-301031
L5b-s-267180
L5b-s-267195
L5b-s-302840
L5ET
L5ET-337966
L5ET-303216
L5ET-302951
L5ET-267033
L5ET-232635
L5ET-267029
L5ET-266839

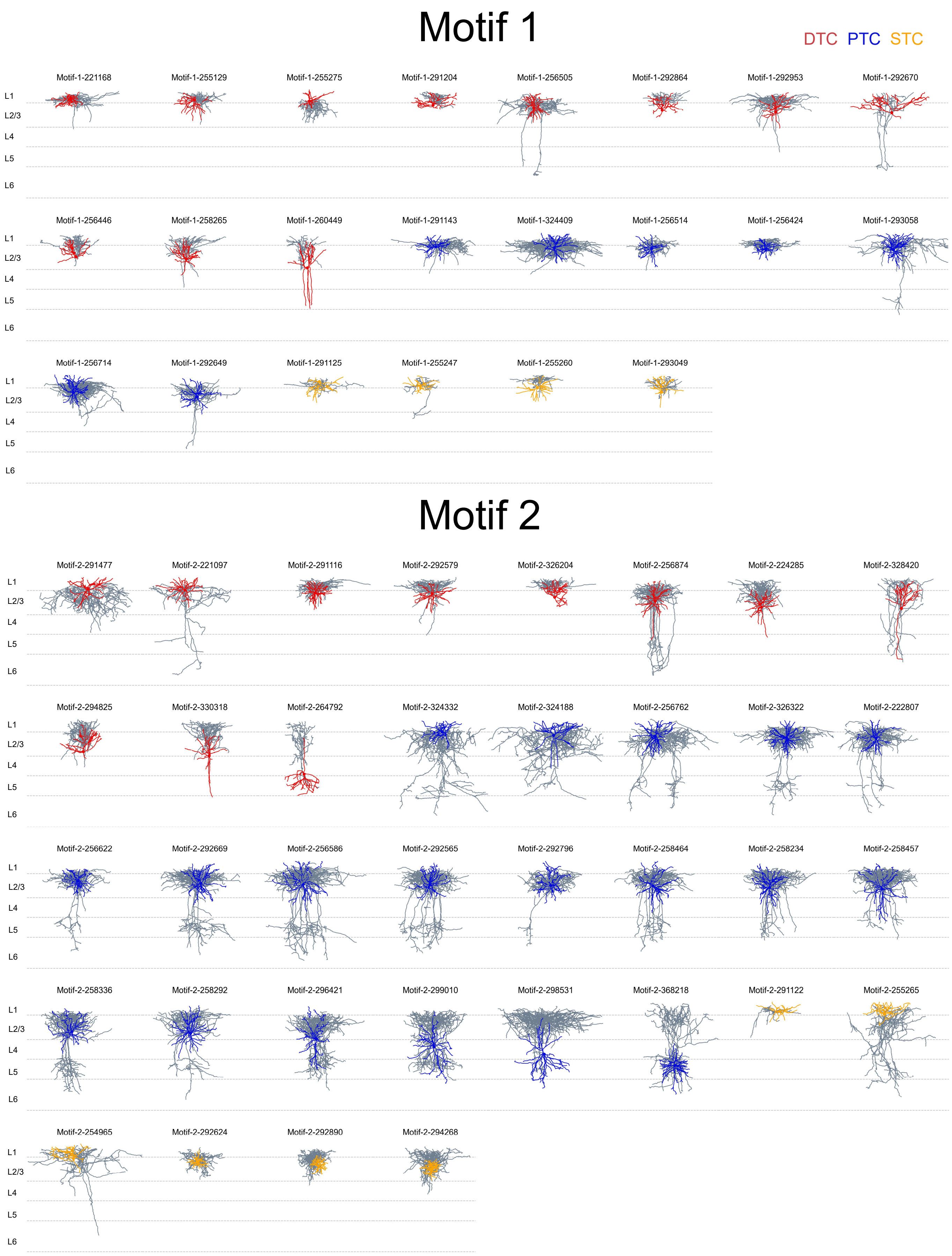
Motif 1
DTC PTC STC
Motif-1-221168
Motif-1-255129
Motif-1-255275
Motif-1-291204
Motif-1-256505
Motif-1-292864
Motif-1-292953
Motif-1-292670
L1
L2/3
L4
L5
L6
Motif-1-256446
Motif-1-258265
Motif-1-260449
Motif-1-291143
Motif-1-324409
Motif-1-256514
Motif-1-256424
Motif-1-293058
L1
L2/3
L4
L5
L6
Motif-1-256714
Motif-1-292649
Motif-1-291125
Motif-1-255247
Motif-1-255260
Motif-1-293049
L1
L2/3
L4
L5
L6
Motif 2
Motif-2-291477
Motif-2-221097
Motif-2-291116
Motif-2-292579
Motif-2-326204
Motif-2-256874
Motif-2-224285
Motif-2-328420
L1
L2/3
L4
L5
L6
Motif-2-294825
Motif-2-330318
Motif-2-264792
Motif-2-324332
Motif-2-324188
Motif-2-256762
Motif-2-326322
Motif-2-222807
L1
L2/3
L4
L5
L6
Motif-2-256622
Motif-2-292669
Motif-2-256586
Motif-2-292565
Motif-2-292796
Motif-2-258464
Motif-2-258234
Motif-2-258457
L1
L2/3
L4
L5
L6
Motif-2-258336
Motif-2-258292
Motif-2-296421
Motif-2-299010
Motif-2-298531
Motif-2-368218
Motif-2-291122
Motif-2-255265
L1
L2/3
L4
L5
L6
Motif-2-254965
Motif-2-292624
Motif-2-292890
Motif-2-294268
L1
L2/3
L4
L5
L6

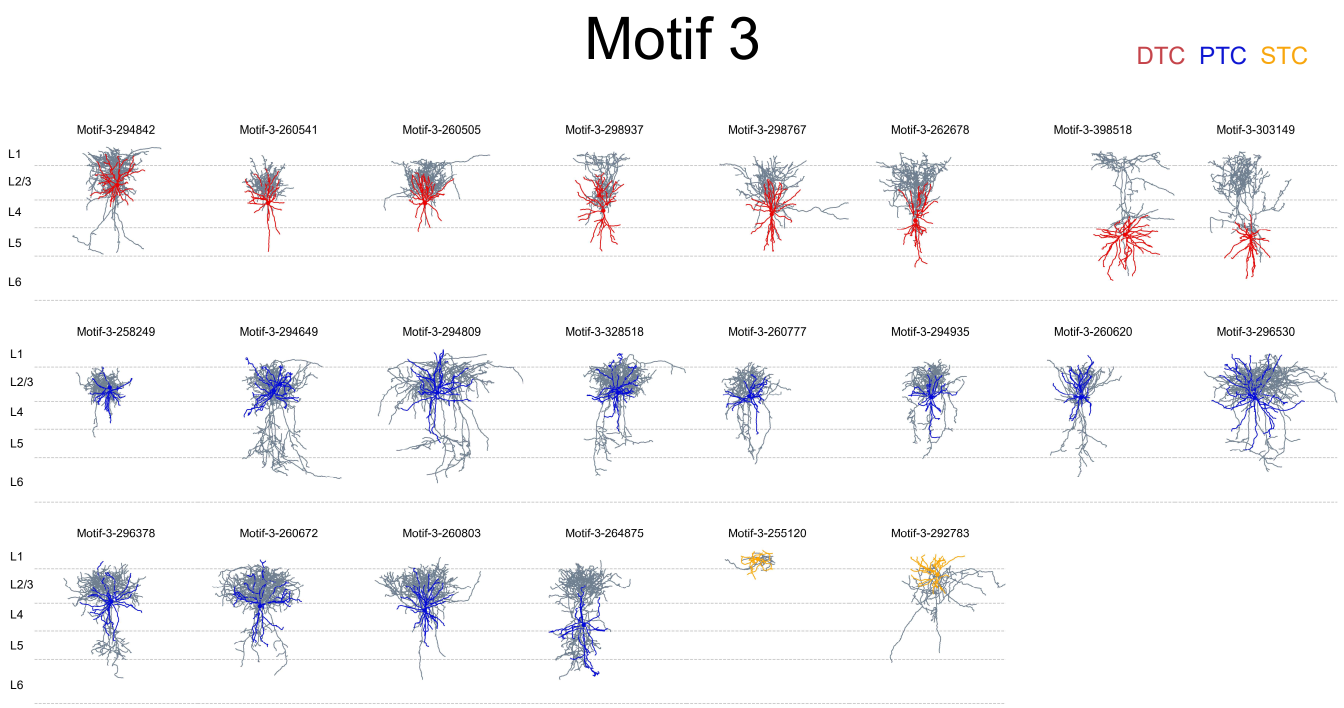
Motif 3
DTC PTC STC
Motif-3-294842
Motif-3-260541
Motif-3-260505
Motif-3-298937
Motif-3-298767
Motif-3-262678
Motif-3-398518
Motif-3-303149
Motif-3-258249
Motif-3-294649
Motif-3-294809
Motif-3-328518
Motif-3-260777
Motif-3-294935
Motif-3-260620
Motif-3-296530
Motif-3-296378
Motif-3-260672
Motif-3-260803
Motif-3-264875
Motif-3-255120
Motif-3-292783
L1
L2/3
L4
L5
L6

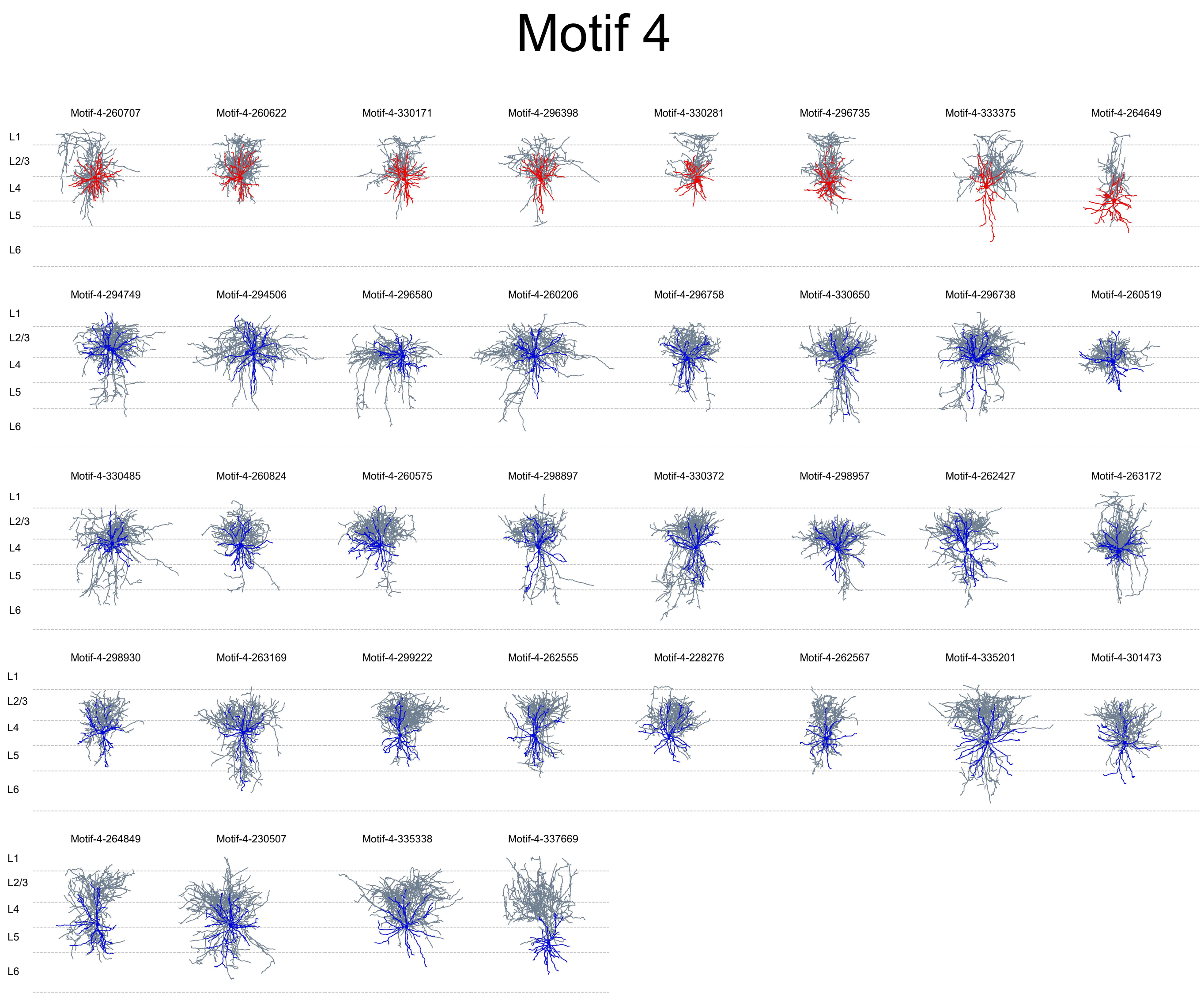
Motif 4
Motif-4-260707
Motif-4-260622
Motif-4-330171
Motif-4-296398
Motif-4-330281
Motif-4-296735
Motif-4-333375
Motif-4-264649
Motif-4-294749
Motif-4-294506
Motif-4-296580
Motif-4-260206
Motif-4-296758
Motif-4-330650
Motif-4-296738
Motif-4-260519
Motif-4-330485
Motif-4-260824
Motif-4-260575
Motif-4-298897
Motif-4-330372
Motif-4-298957
Motif-4-262427
Motif-4-263172
Motif-4-298930
Motif-4-263169
Motif-4-299222
Motif-4-262555
Motif-4-228276
Motif-4-262567
Motif-4-335201
Motif-4-301473
Motif-4-264849
Motif-4-230507
Motif-4-335338
Motif-4-337669
L1
L2/3
L4
L5
L6

### Motif 5

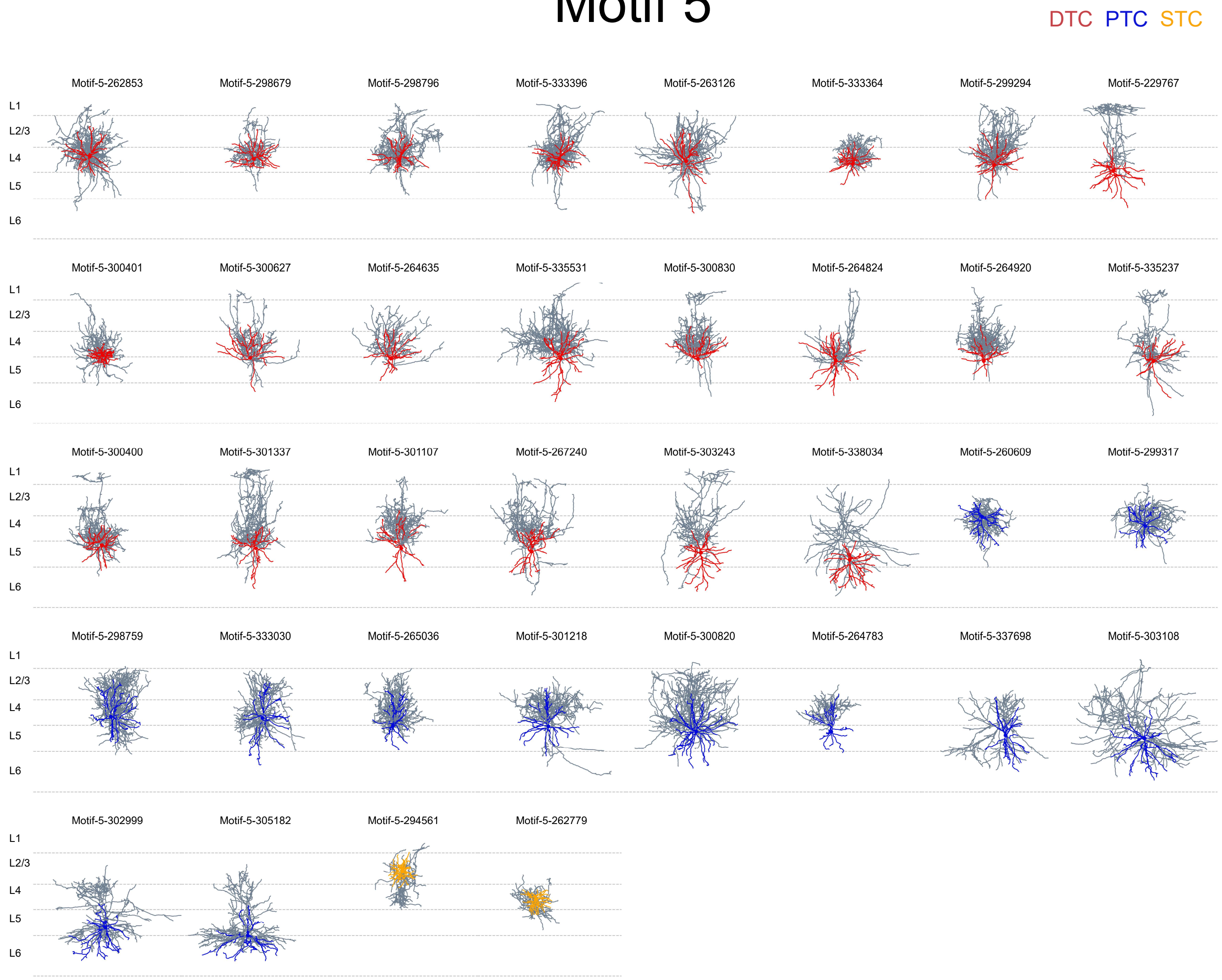


### Motif 6

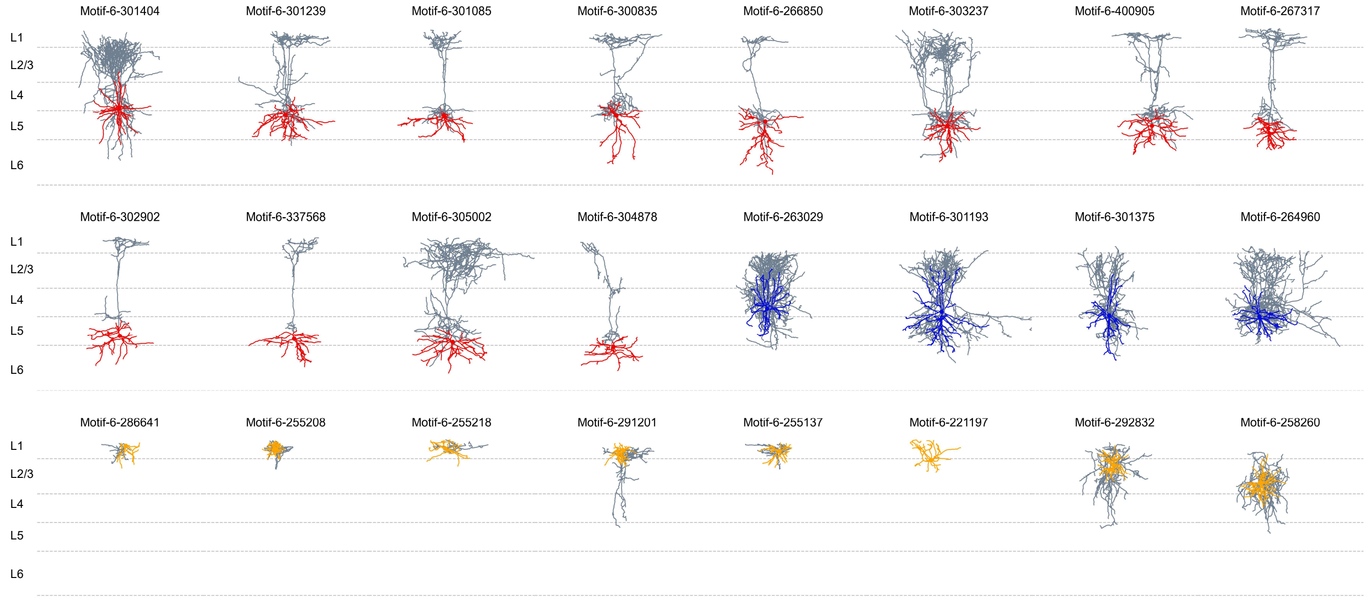

### Motif 7

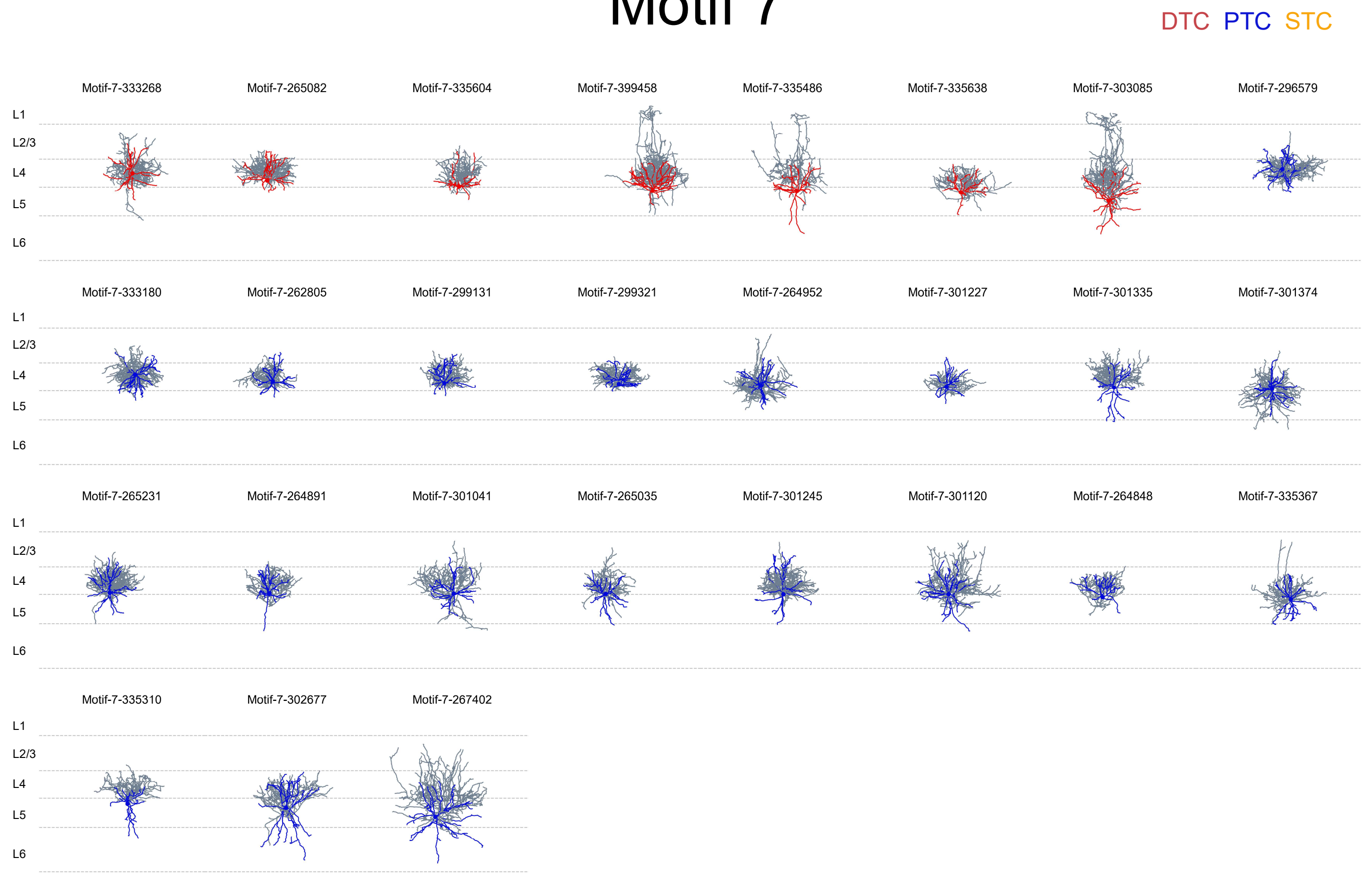


### Motif 8/10

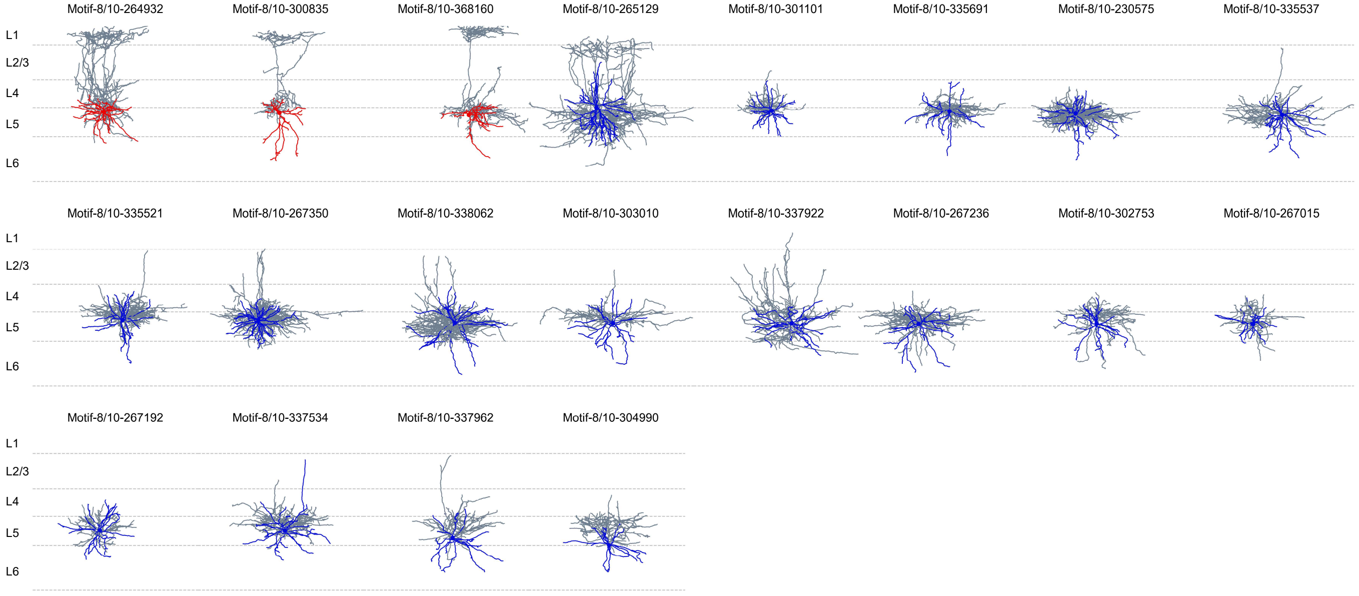

### Motif 9a

DTC PTC STC

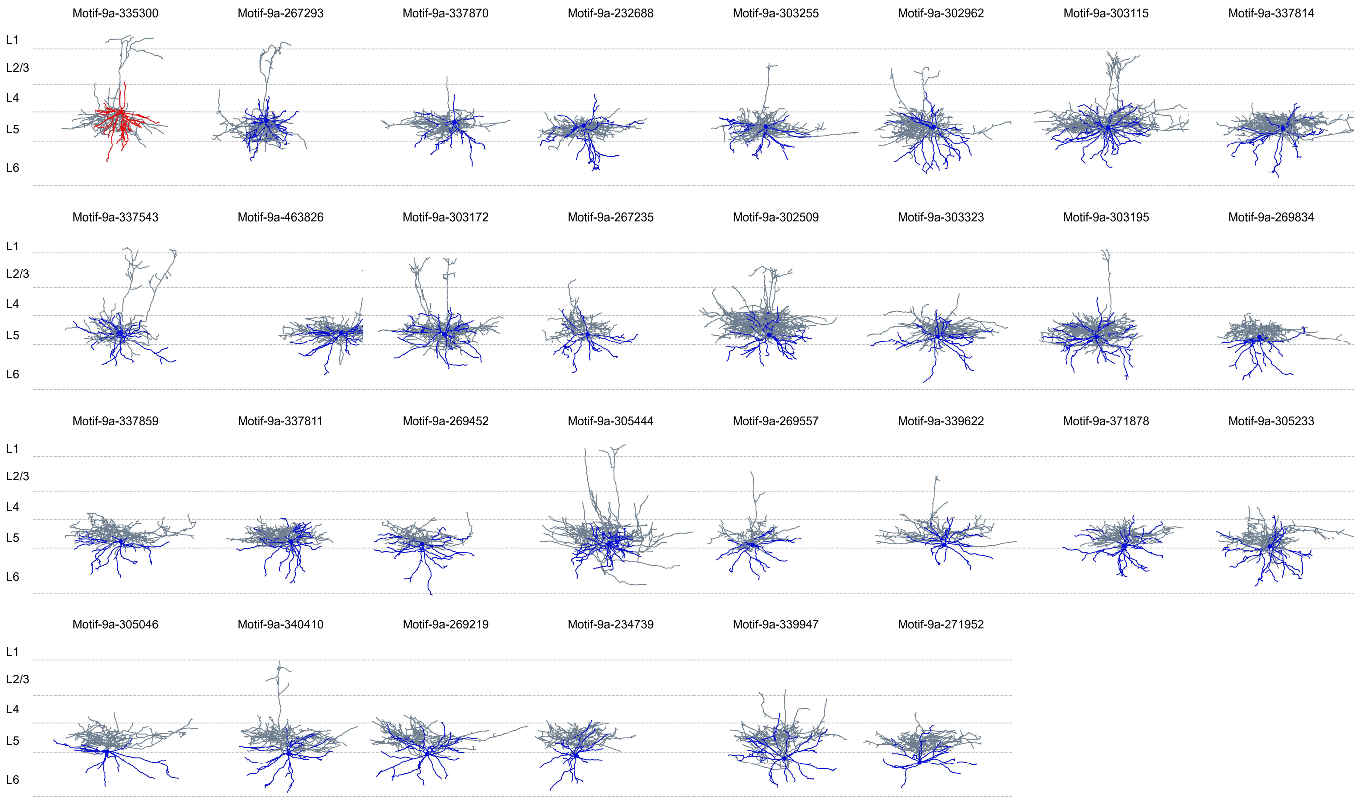


### Motif 9b

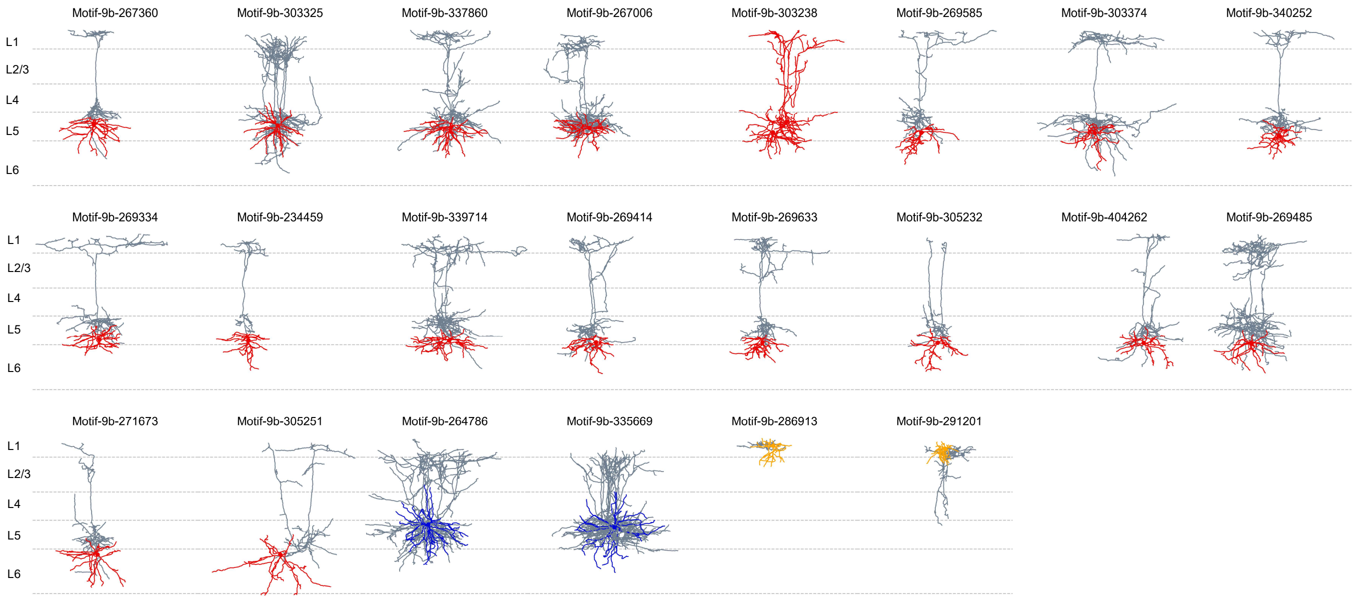

### Motif 11/12

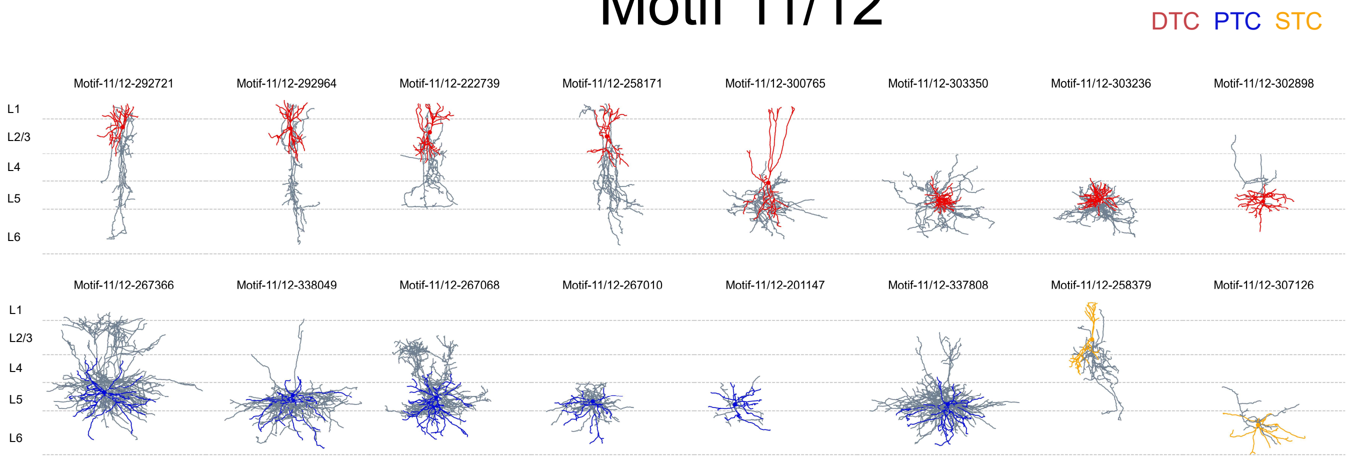


### Motif 13

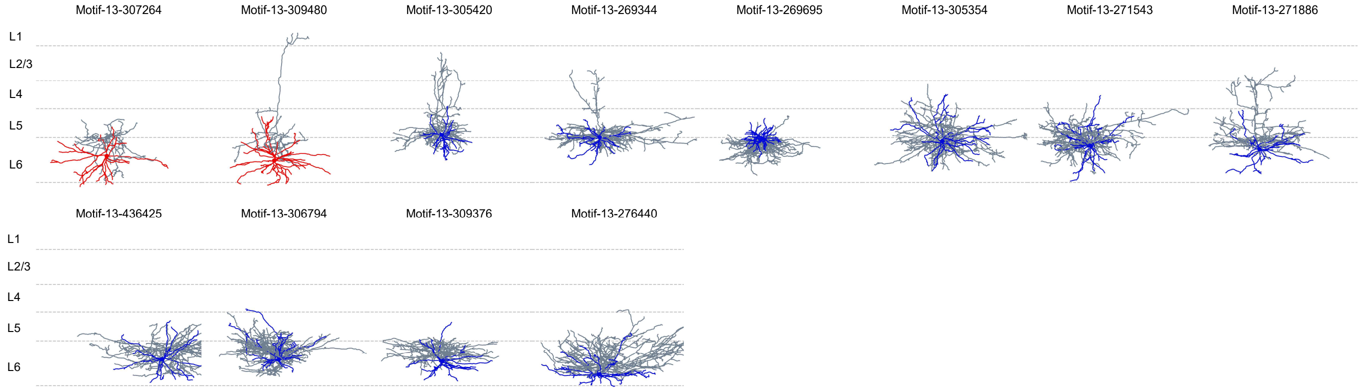


### Motif 14

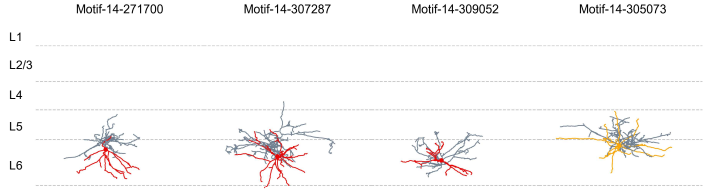


### Motif 15/16

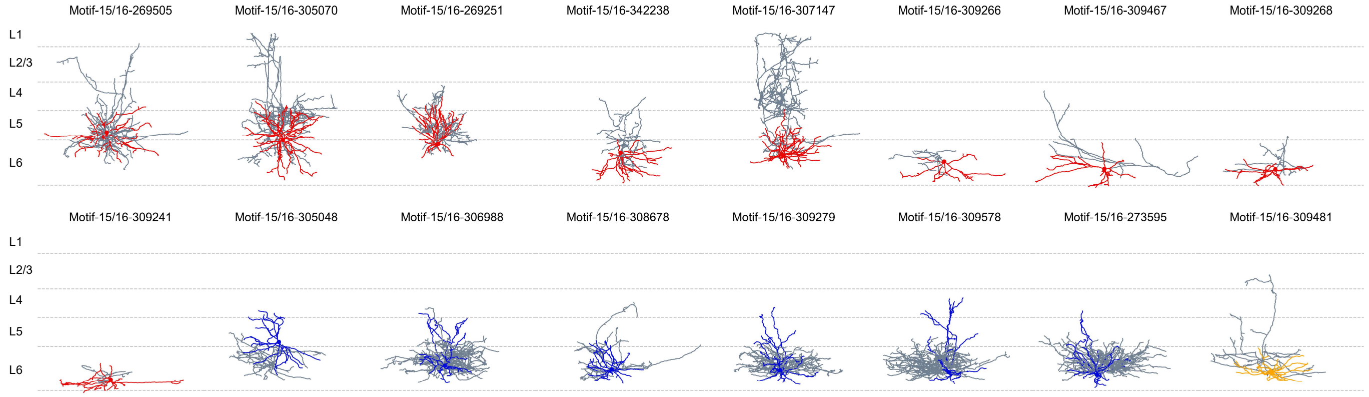

### Motif 17/18

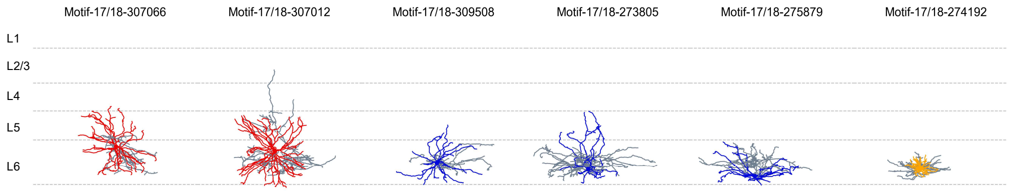
Motif-17/18-307066
Motif-17/18-307012
Motif-17/18-309508
Motif-17/18-273805
Motif-17/18-275879
Motif-17/18-274192
L1
L2/3
L4
L5
L6
